# Unexpected inhibition of the lipid kinase PIKfyve reveals an epistatic role for p38 MAPKs in endolysosomal fission and volume control

**DOI:** 10.1101/2023.03.13.532495

**Authors:** Daric J. Wible, Zalak Parikh, Eun Jeong Cho, Miao-Der Chen, Somshuvra Mukhopadhyay, Kevin N. Dalby, Shankar Varadarajan, Shawn B. Bratton

**Affiliations:** The University of Texas MD Anderson Cancer Center, Department of Epigenetics & Molecular Carcinogenesis, Houston, Texas 77054; The University of Texas at Austin, Targeted Therapeutic Drug Discovery and Development Program, Division of Chemical Biology & Medicinal Chemistry, College of Pharmacy, Austin, Texas 78712; The University of Texas at Austin, Division of Pharmacology & Toxicology, College of Pharmacy, Austin, Texas 78712; University of Liverpool, Institute of Translational Medicine, Liverpool L69 3BX, United Kingdom

**Keywords:** p38 MAPK, Rab7, ARF1, PIKfyve, lysosome fission

## Abstract

p38 mitogen-activated protein kinases (MAPKs) regulate early endocytic trafficking, but their effects on late endocytic trafficking remain unclear. Herein, we report that the pyridinyl imidazole p38 MAPK inhibitors, SB203580 and SB202190, induce a rapid but reversible Rab7-dependent accumulation of large cytoplasmic vacuoles. While SB203580 did not induce canonical autophagy, phosphatidylinositol 3-phosphate [PI(3)P] accumulated on vacuole membranes, and inhibition of the class III PI3-kinase (PIK3C3/VPS34) suppressed vacuolation. Ultimately, vacuolation resulted from the fusion of ER/Golgi-derived membrane vesicles with late endosomes and lysosomes (LELs), combined with an osmotic imbalance in LELs that led to severe swelling and a decrease in LEL fission. Since PIKfyve inhibitors induce a similar phenotype by preventing the conversion of PI(3)P to PI(3,5)P2, we performed *in vitro* kinase assays and found that PIKfyve activity was unexpectedly inhibited by SB203580 and SB202190, corresponding to losses in endogenous PI(3,5)P2 levels in treated cells. However, vacuolation was not entirely due to ‘off-target’ inhibition of PIKfyve by SB203580, as a drug-resistant p38α mutant suppressed vacuolation. Moreover, genetic deletion of both p38α and p38β rendered cells dramatically more sensitive to PIKfyve inhibitors, including YM201636 and apilimod. In subsequent ‘washout’ experiments, the rate of vacuole dissolution upon the removal of apilimod was also significantly reduced in cells treated with BIRB-796, a structurally unrelated p38 MAPK inhibitor. Thus, p38 MAPKs act epistatically to PIKfyve to promote LEL fission; and pyridinyl imidazole p38 MAPK inhibitors induce cytoplasmic vacuolation through the combined inhibition of both PIKfyve and p38 MAPKs.

## INTRODUCTION

p38 MAPKs regulate a plethora of biological responses through phosphorylation of key cellular protein substrates. During early endocytosis, for example, p38 MAPKs increase the rates of receptor internalization mediated by the Rab GTPase, Rab5, through phosphorylation and activation of GDP dissociation inhibitor α (GDIα) and early endosome antigen 1 (EEA1) (1–4). Phosphorylation of EEA1, in particular, promotes its removal from early endosomes and inhibits homotypic fusion (2). Once formed, the maturation of early endosomes into late endosomes (LEs) is marked by the replacement of Rab5 with Rab7 (5). The mechanisms that mediate the transfer of LE contents to lysosomes remain somewhat controversial, but LEs are thought to directly fuse with lysosomes to produce endo-lysosomal “hybrid” organelles (6). Following digestion of cargo within hybrid organelles, LEs and lysosomes are thought to be recovered as a result of fission (6). As noted above, while p38 MAPKs clearly regulate early endocytic events, it remains unclear whether they also regulate late endocytic events, including the reformation of LEs and lysosomes.

Similar to endocytosis, ADP ribosylation factor (ARF) GTPase family members, secretion-associated and *ras* superfamily-related gene (SAR1) and ARF1, regulate the transport of protein-containing vesicles between the endoplasmic reticulum (ER) and Golgi compartments (7). ER proteins are exported from “ER exit sites” (ERES) following recruitment and activation of the SAR1/coat protein II (COPII) complex, which in turn mediates the budding of vesicles or tubules from the ER (8). The ARF1/COPI complex, in addition to regulating retrograde transport, mediates the differentiation and detachment of vesicles from the ER and propels them towards the center of the cell, where they form Golgi (9). Notably, disruption of protein export from “ER exit sites” (ERES) induces ER stress, which typically activates both c-Jun N-terminal kinases (JNKs) and p38 MAPKs (10–12).

During autophagy, isolation membranes are thought to originate largely from ER or Golgi membranes through a process involving the essential autophagy protein ATG9A, a lipid scramblase that is normally incorporated into vesicles that emanate from the *trans*-Golgi network (TGN) (13,14). These vesicles are recruited to pre-autophagosomal structures (PAS) at ERES, where they serve as the initial membrane source for the growth of phagophores (15,16); and p38 MAPKs reportedly regulate ATG9A and autophagy through phospho-dependent sequestration of p38IP, an ATG9A interactor that influences its trafficking (17). Importantly, the role(s) for p38 MAPKs in promoting autophagy appeared to be at odds with earlier reports, which suggested that the p38 MAPK inhibitors, SB203580 and SB202190, induced cell death in some cancer cells through unrestrained autophagy (18,19). Subsequent reports by others, however, argued that cytoplasmic vacuolation induced by both drugs was unrelated to p38 MAPK inhibition (20–22), once again raising doubts as to the true roles of p38 MAPKs in autophagy, since the same inhibitors are utilized in most studies.

Herein, using a litany of approaches, we demonstrate that pyridinyl imidazole p38 MAPK inhibitors induce profound cytoplasmic vacuolation independently of either endocytosis or autophagy, both of which are suppressed in cells. Growth of these vacuoles relies upon Rab7-dependent membrane fusion and the contribution of ER/Golgi-derived membrane vesicles (and pre-existing autophagosomes) that likely emanate from ERES. Most surprisingly, SB203580 and SB202190 directly inhibit the lipid kinase PIKfyve (also known as PIP5K3), which normally resides on endo-lysosomal membranes, catalyzes the conversion of PI(3)P to PI(3,5)P2, and in turn, positively regulates the activities of several ion channels. While PIKfyve inhibition is sufficient to trigger an osmotic disturbance in LELs and swelling, simultaneous inhibition of p38 MAPKs induces even more profound vacuolation, apparently through a concomitant block in fission. Thus, in summary, using a phenotypic drug discovery approach to identify the “off-target(s)” of SB203580 and SB202190 that are responsible for vacuolation, we have identified a previously unrecognized role for p38 MAPKs in regulating endo-lysosomal fission that is epistatic to PIKfyve.

## RESULTS

### Pyridinyl imidazole p38 MAPK inhibitors induce cytoplasmic vacuolation

In studies begun many years ago, we observed that the p38 MAPK inhibitors, SB203580 and SB202190, induced profound cytoplasmic vacuolation in multiple cancer cell lines including prostate (DU145, PC-3), lung (A549), colon (HCT116, HT-29), and cervical cancers (HeLa), as well as immortalized mouse embryonic fibroblasts (MEFs) (Fig. 1A,B; fig. S1A). In marked contrast, pharmacological inhibitors of other stress and growth-related kinases, including c-Jun N-terminal kinases (JNKs, SP600125), extracellular signal-regulated kinases (ERKs, PD98059), mammalian target of rapamycin (mTOR, rapamycin), and phosphoinositide 3-kinases (PI3Ks, LY294002) failed to induce comparable levels of vacuolation (fig. S1B). SB203580-induced vacuolation did not lead to a substantial increase in cell death across multiple cell types, at least over a 24 h time span (Fig. 1B), and caused only a moderate reduction in DU145 cell proliferation during several days in culture, as determined by CFDA labeling (Fig. 1C). Vacuolation was also entirely reversible with most cells returning to normal within 8 h following the washout of drug (Fig. 1D; fig. S1C).

**Figure 1.**
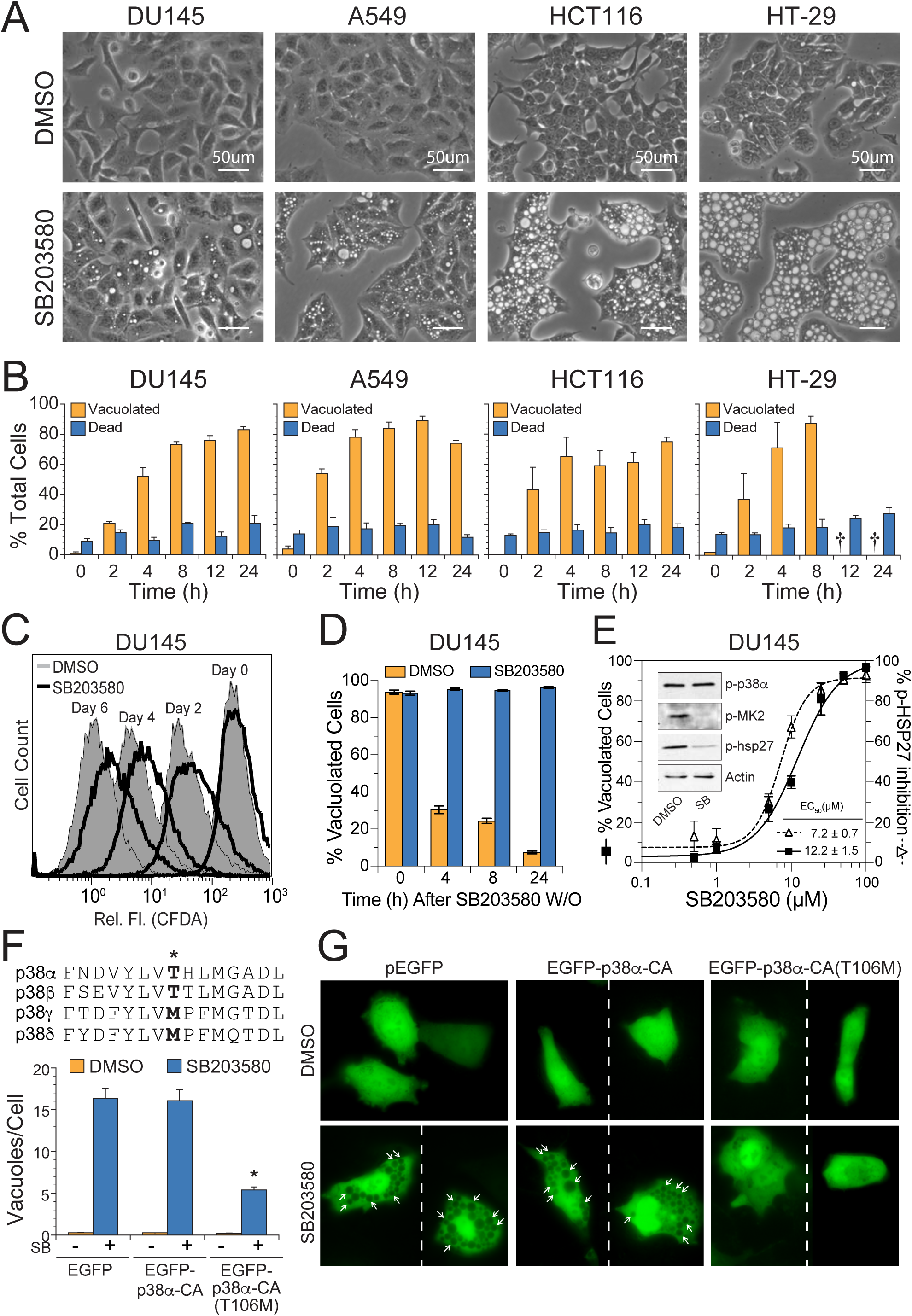
Pyridinyl imidazole p38 MAPK inhibitors induce cytoplasmic vacuolation. *(A and B)* Various transformed and cancer cell lines were treated with DMSO (control) or SB203580 (50 µM) for 0-24 h and assessed for vacuolation and cell death by phase-contrast microscopy and flow cytometry (annexin V-propidium iodide staining), respectively. *(C)* DU145 cells were stained with CFDA-SE, exposed to DMSO (shaded peaks) or SB203580 (empty peaks), and monitored for changes in cell proliferation, as described in the methods. *(D)* DU145 cells were treated with SB203580 (50 µM) for 24 h and then washed and replaced with fresh media ± SB203580 for 4-24 h. At each time point following the washout, cells were examined for vacuolation by phase-contrast microscopy (*see* also Fig. S1C). *(E)* DU145 prostate cancer cells were treated with increasing concentrations of the pharmacological p38 MAPK inhibitor, SB203580 (0 – 100 µM), for 24 h and examined for signs of vacuolation by phase-contrast microscopy (200X). *Inset:* SB203580 (50 µM) inhibited p38-dependent sequential phosphorylation of MK2 and HSP27. Concentration-dependent inhibition of HSP27 phosphorylation by SB203580 (0 – 100 µM) was also determined by western blotting (*see* Fig. S1D) with individual bands scanned, quantified with ImageJ software, and plotted as the percent of p-HSP27 inhibited. *(F and G)* DU145 cells were transiently transfected with expression plasmids encoding EGFP, constitutively active (D176A/F327S) p38α (EGFP-p38α-CA), or p38α-CA containing an additional mutation to the gatekeeper residue (T106M) that renders p38α resistant to SB203580 [EGFP-p38α-CA (T106M)]. EGFP-positive cells were then evaluated by fluorescence microscopy for the number of vacuoles present per cell.

SB203580 inhibited the p38 MAPK pathway, as indicated by a concentration-dependent decrease in the sequential phosphorylation of MAP kinase-activated protein kinase 2 (MK2) and heat shock protein 27 (HSP27) (Fig. 1E; fig. S1D), and pathway inhibition correlated strongly with an increase in cellular vacuolation (Fig. 1E; fig. S1E, R^2^ = 0.82). SB203580 inhibits the activities of p38α and p38β isoforms, but not p38δ and p38γ, due to the presence of a threonine (rather than a methionine) “gatekeeper” residue in their ATP-binding pockets (Fig. 1F, inset) (23,24). Therefore, to confirm that selective inhibition of p38α/β MAPKs by SB203580 was responsible for (or at least contributed to) cytoplasmic vacuolation, we performed rescue experiments wherein cells were transfected with an EGFP-tagged, constitutively active mutant of p38α (D176A/F327S; termed p38α-CA) (25), possessing either the wild-type threonine or a mutated methionine residue [p38α-CA(T106M)] (26). Upon exposure of cells to SB203580, those transfected with either EGFP or EGFP-p38α-CA underwent normal vacuolation, whereas those transfected with EGFP-p38α-CA(T106M) were resistant to SB203580-induced vacuolation (Fig. 1F,G). Thus, selective inhibition of p38α/β MAPKs (hereafter referred to simply as p38 MAPKs) by SB203580 appeared to be responsible for the observed vacuolation.

### SB203580 does not induce autophagy but requires PIK3C3 activity to induce vacuolation

A previous report suggested that vacuolation of SB202190-treated cells resulted from autophagic cell death that was otherwise suppressed by p38 MAPKs (18,19). While SB203580 induced very little cell death in our hands (Fig. 1B), inhibition of the autophagy-essential class III PI3-kinase PIK3C3 (also known as VPS34) with 3-methyladenine (3-MA) (27) did dramatically inhibit vacuolation in all cell lines tested (Fig. 2A), as did stable shRNA knockdown of the PIK3C3 accessory protein/regulator Beclin 1 in DU145 cells (Fig. 2B). To verify that 3-MA effectively inhibited PIK3C3 activity, we generated an EGFP construct fused to a single PX domain of p40Phox (PX-EGFP), as well as a PI(3)P-binding mutant (PX-R57Q-EGFP) (28). Following exposure to SB203580, many of the small to medium-sized vacuoles were clearly labeled with PX-EGFP but not with PX-R57Q-EGFP (Fig. 2C, *images ii* and *iv*). Cotreatment with 3-MA or knockdown of Beclin 1, on the other hand, inhibited the formation of PI(3)P and produced a diffuse cytoplasmic staining pattern, similar to that observed in control cells (Fig. 2C, *images i*, *iii*, and *vi*). Thus, SB203580 induced the formation of cytoplasmic vacuoles, many of which were enriched in PI(3)P; and inhibition of PIK3C3 with 3-MA or knockdown of Beclin 1 inhibited both PI(3)P production and vacuole formation.

**Figure 2.**
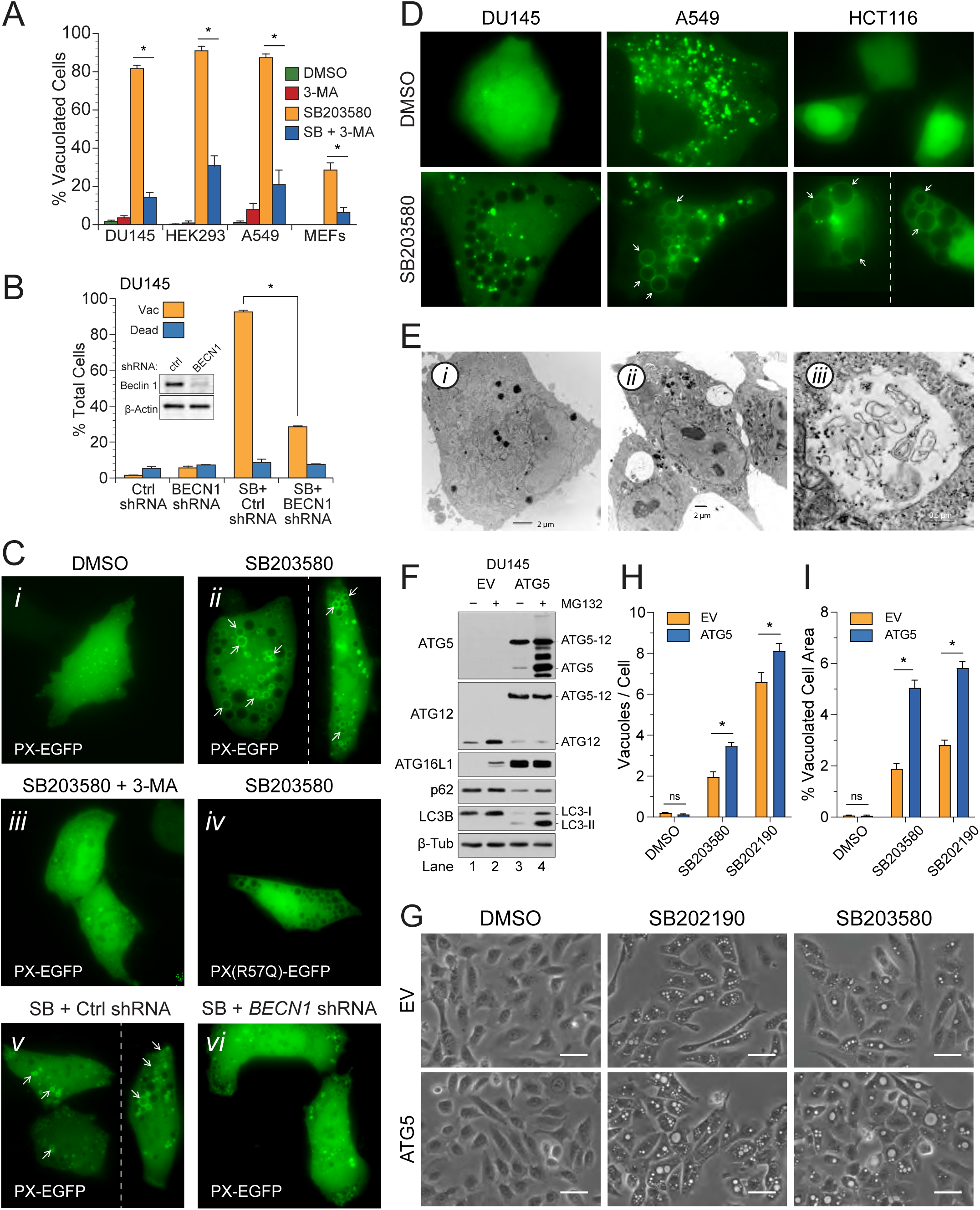
SB203580 does not stimulate autophagy but does induce vacuolation in a PIK3C3-dependent manner and incorporates pre-existing autophagosome membranes. *(A)* DU145, HEK-293, A549, and MEFs were exposed to SB203580 (50 µM) ± the class III PI3 kinase inhibitor, 3-MA (5 mM), and examined for vacuolation. For each cell-type, asterisks indicate that 3-MA was significantly different from the SB treatment alone (one-way ANOVA with Student-Neumann-Keuls posthoc analysis; p<0.05). *(B)* DU145 cells were stably transfected with scrambled control or *beclin 1* shRNA plasmids, treated with DMSO (control) or SB203580 (50 µM), and examined for vacuolation and cell death. *(Inset)* Beclin 1 expression levels were determined by immunoblotting. *(C)* DU145 cells were transiently transfected with the PI(3)P probe, PX-EGFP, or the binding mutant R57Q, and subsequently treated with *(i)* DMSO or *(ii-iv)* SB203580 ± 3-MA for 24 h. Similarly, (*v*) control and (*vi*) Beclin 1-depleted cells were also transfected with PX-EGFP and treated with SB203580 (50 µM) for 24 h and examined by fluorescence microscopy. *(D)* DU145, A549, and HCT116 cancer cell lines were transiently transfected with EGFP-LC3 and treated with SB203580 (50 µM) for 24 h. Some but not all LC3-labeled vacuoles are indicated with arrows (*see* also Fig. S2A). *(E)* DU145 cells were treated with *(i)* DMSO or *(ii-iii)* SB203580 (50 µM) for 24 h and analyzed by transmission electron microscopy. Whole cells in panels *i* and *ii* were magnified 4,400X and 2,800X, respectively (bars = 2 µm); and both large translucent and partially-filled vacuoles (panel *ii*) were visible in SB203580-treated cells. A representative vacuole (panel *iii*) was magnified 28,000X (bar = 500 nm) and clearly contained partially digested material. *(F-I)* DU145 were reconstituted with ATG5 to restore basal autophagy; exposed to SB203580 (50 µM) or SB202190 (50 µM) for 24 h; and evaluated by phase-contrast microscopy for vacuolation.

In parallel experiments, to confirm the formation of autophagosomes and determine if the large vacuoles arose out of an increase in autophagy, we transfected cells with EGFP-tagged microtubule-associated protein 1 light chain 3 (EGFP-LC3) and examined them for evidence of autophagosome formation, as indicated by the accumulation of EGFP-LC3 puncta (29). No puncta were observed in DU145, HCT116, or HEK293 control cells, whereas numerous puncta were observed in A549, HELA, and MEF control cells, suggesting that the latter possessed higher rates of basal autophagy (Fig. 2D and S2A, upper panels). Following treatment with SB203580, vacuoles in all cell lines were decorated with EGFP-LC3 with the notable exception of DU145 cells (Fig. 2D and S2A, lower panels). Cell lines that displayed the highest levels of basal autophagy, prior to treatment with SB203580, likewise exhibited the greatest degree of vacuole labeling following treatment, implying that EGFP-LC3 from pre-existing autophagosomes was incorporated into developing vacuoles following fusion (Fig. 2D and S2A).

We were puzzled at the time, however, as to why the SB203580-induced vacuoles in DU145 cells were not labeled with EGFP-LC3 (Fig. 2D, lower panel), so we examined DU145 cells using transmission electron microscopy (TEM). To our surprise, all SB203580-induced vacuoles were found to be single-bilayer rather than the double-bilayer membrane structures characteristic of autophagosomes (Fig 2E, fig. S2B). Larger vacuoles (> 1 μm diameter) were mostly empty (Fig. 2E, *image ii*), whereas smaller vacuoles, which were not readily identifiable by phase-contrast microscopy, contained undigested vesicles and cytoplasmic material reminiscent of multivesicular bodies (MVBs) (Fig. 2E, *image iii*; fig. S2B, *images iv*-*vi*). We would later discover that DU145 cells were naturally deficient in ATG5, due to a donor splice site gene mutation, and thus were incapable of undergoing canonical autophagy (Fig. 2F, lanes 1 and 2) (30). Long-lived protein degradation (LLPD) assays confirmed an absence of autophagy in DU145 cells in response to SB203580 or starvation (fig. S2C). Notably, however, SB203580 also failed to stimulate an increase in LLPD in “autophagy-competent” A549, HCT116, or HT-29 cells (fig. S2C) and stimulated the formation of empty single-bilayer vacuoles and enlarged MVBs or amphisomes in HCT116 cells, as determined by TEM (fig. S2D).

Even though SB203580 did not induce autophagy (Fig. 2D,E; fig. S2B-D), the fact that EGFP-LC3 readily labeled vacuoles in autophagy-competent cell types (Fig. 2D; fig. S2A) led us to question whether autophagosomes, resulting from basal autophagy, might still contribute membrane to the expanding vacuole. To evaluate this possibility, we reconstituted DU145 cells with ATG5 (Fig. 2F, lanes 3 and 4) and found that restoration of autophagy slightly increased the number of vacuoles formed per cell, following treatment with SB203580 (Fig. 2G,H), but more substantially increased the size of those vacuoles (Fig. 2G,I). Thus, in summary, SB203580 stimulated vacuolation in a PIK3C3-dependent manner but did not induce canonical autophagy *per se*, although preexisting autophagosomes, resulting from basal autophagy, could contribute to vacuole expansion.

### SB203580 stimulates Rab7-dependent enlargement of MVBs/late endosomes and lysosomes

Two distinct PIK3C3 complexes selectively associate with phagophores during autophagy or endosomes during endocytosis, depending upon their interactions with adapter proteins, ATG14L or UVRAG, respectively (31,32). Given that PIK3C3 inhibition prevented SB203580-induced vacuolation, even in autophagy-deficient DU145 cells (Fig. 2A,B), we considered the possibility that dysregulated endocytosis might be responsible for vacuolation. Indeed, PIK3C3 regulates endocytosis following early endosome recruitment and activation by GTP-bound Rab5 (33), and previous reports indicate that p38 MAPKs can phosphorylate a number of factors associated with Rab5^+^ early endosomes, including guanyl-nucleotide dissociation inhibitor (GDI, Ser-121) and early endosome antigen 1 (EEA1, Thr-1392) (1,2). However, in most cases, p38 MAPK activity promotes endocytosis, and consistent with these earlier studies, we found that SB203580 did not stimulate the labeling of vacuoles with either EGFP-tagged Rab5 or EEA1 (Fig. 3A; fig. S3A, upper panels). More importantly, overexpression of a GDP-locked dominant-negative Rab5(N133I) failed to inhibit vacuolation (Fig. 3B,C), as did overexpression of functional phosphomimetics of EEA1(T1392E) and GDI(S121E) (fig. S3, A-C) (1,2). Overexpression of AP180C, an inhibitor of clathrin-mediated endocytosis, likewise failed to inhibit vacuolation (data not shown). Finally, we found that SB203580 actually inhibited the internalization of fluid-phase markers, Dextran-FITC and BSA-FITC, ruling out a significant role for macropinocytosis in vacuolation as well (fig. S3D,E). Thus, inhibition of p38 MAPKs did not appear to induce cytoplasmic vacuolation through a *de novo* increase in endocytosis, homotypic fusion of early endosomes, or macropinocytosis.

**Figure 3.**
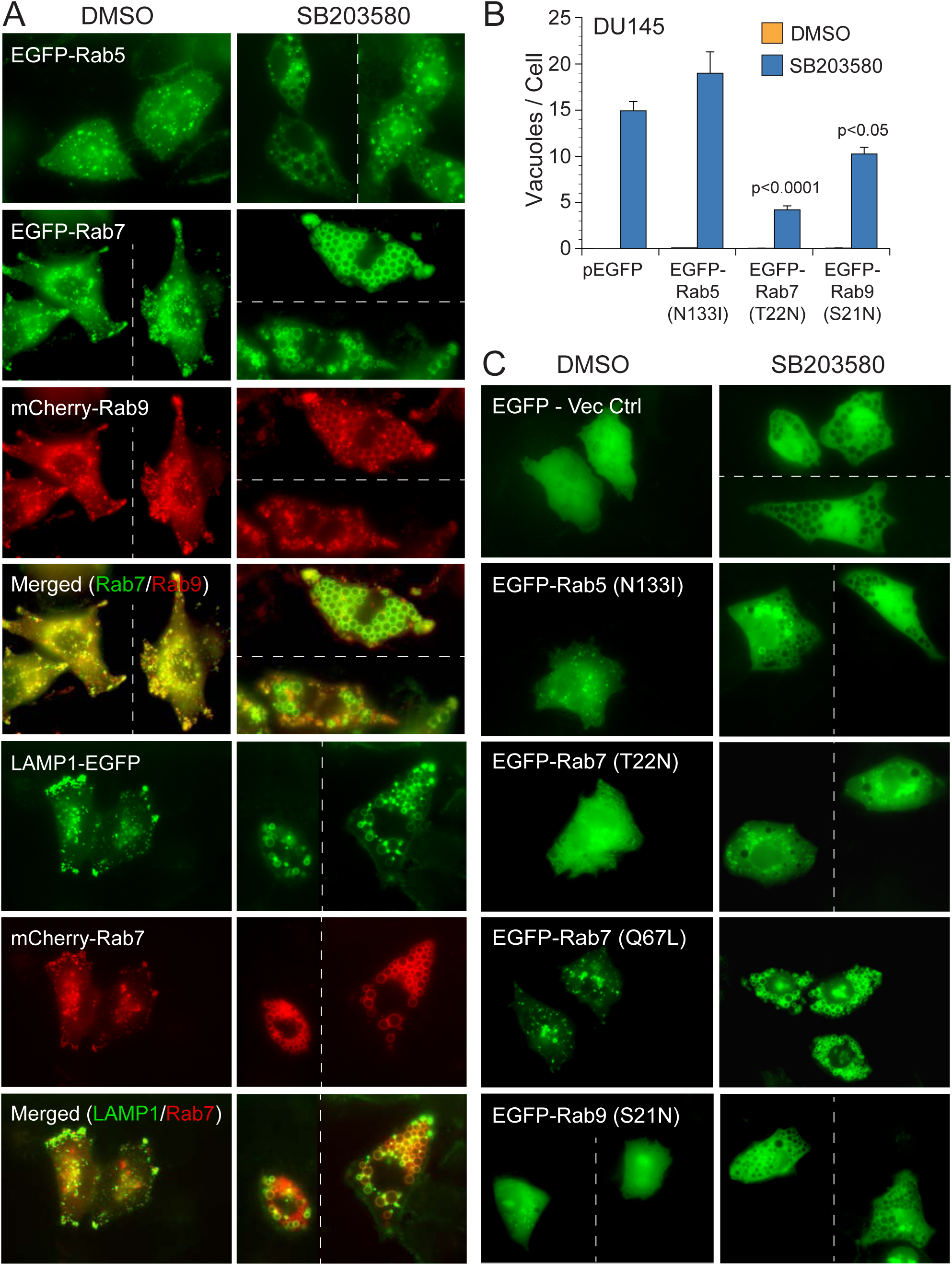
SB203580-induced vacuolation results from Rab7-dependent fusion of LEs and lysosomes. *(A)* DU145 cells were cotransfected with various combinations of EGFP or mCherry-labeled Rab5, Rab7, Rab9, and/or LAMP1. The cells were then treated with DMSO (control) or SB203580 (50 µM) for 24 h and examined for colocalization with the vacuoles and/or one another. *(B and C)* DU145 cells were similarly transfected with either EGFP (empty vector), constitutively active EGFP-tagged Rab7(Q67L), or dominant-negative Rab5(N133I), Rab7(T22N), or Rab9(S21N) mutants. Cells were then exposed to DMSO or SB203580 (50 µM), and the number of vacuoles was counted in at least 50 cells by fluorescence microscopy. Each experiment was performed in triplicate, and each data point represents mean ± SEM.

To determine if SB203580-induced vacuoles were instead labeled with markers for MVBs/late endosomes (LEs) or lysosomes, we transfected DU145 cells with fluorescent fusion proteins of Rab7, Rab9, and LAMP1, alone or in combination. All vacuoles were extensively decorated with Rab7 and a fraction co-labeled with Rab9 and/or LAMP1 (Fig. 3A; fig. S3F). More importantly, vacuolation was substantially reduced following the expression of a GDP-locked dominant-negative mutant of Rab7(T22N) and, to a much lesser extent, Rab9(S21N) (Fig. 3B,C). Conversely, vacuolation was dramatically enhanced by a constitutively-active mutant of Rab7(Q67L), so much so that accurately counting vacuoles was impractical (Fig. 3C). Collectively, our results strongly indicated that Rab7 was required for vacuolation, suggesting that the vacuoles induced by SB203580 were enlarged LEs and lysosomes (hereafter referred to as LELs), consistent with our previous TEM results (Fig. 2E; fig. S2B,D).

### ER/Golgi membranes and ATG9A support expansion of SB203580-induced vacuoles

In order to determine the source of membranes contributing to vacuole expansion, we next turned our attention to ER/Golgi membranes, where Beclin 1 is partially localized (34) and PIK3C3 often catalyzes the formation of phagophores (35,36). We speculated that membranes emanating from the ER/Golgi, be they single membrane phagophores that failed to undergo closure in ATG5-deficient cells or fully mature autophagosomes, might similarly contribute to vacuolation through Rab7-mediated fusion with LELs (Fig. 4A). First, to visualize ER membranes, we transfected wild-type DU145 cells with a construct that targeted EGFP selectively to ER membranes (EGFP-Cytochrome [Cyt.] b5) (37). Following SB203580 treatment, EGFP-Cyt. b5-associated membranes appeared to fully surround some vacuoles (fig. S4A, *white arrows*) while only partially surrounding others (fig. S4A, *red arrows*), which we speculated might be “omegasomes” where ER exit sites (ERES) and pre-autophagosomal structures (PAS) are known to reside (35,38,39).

**Figure 4.**
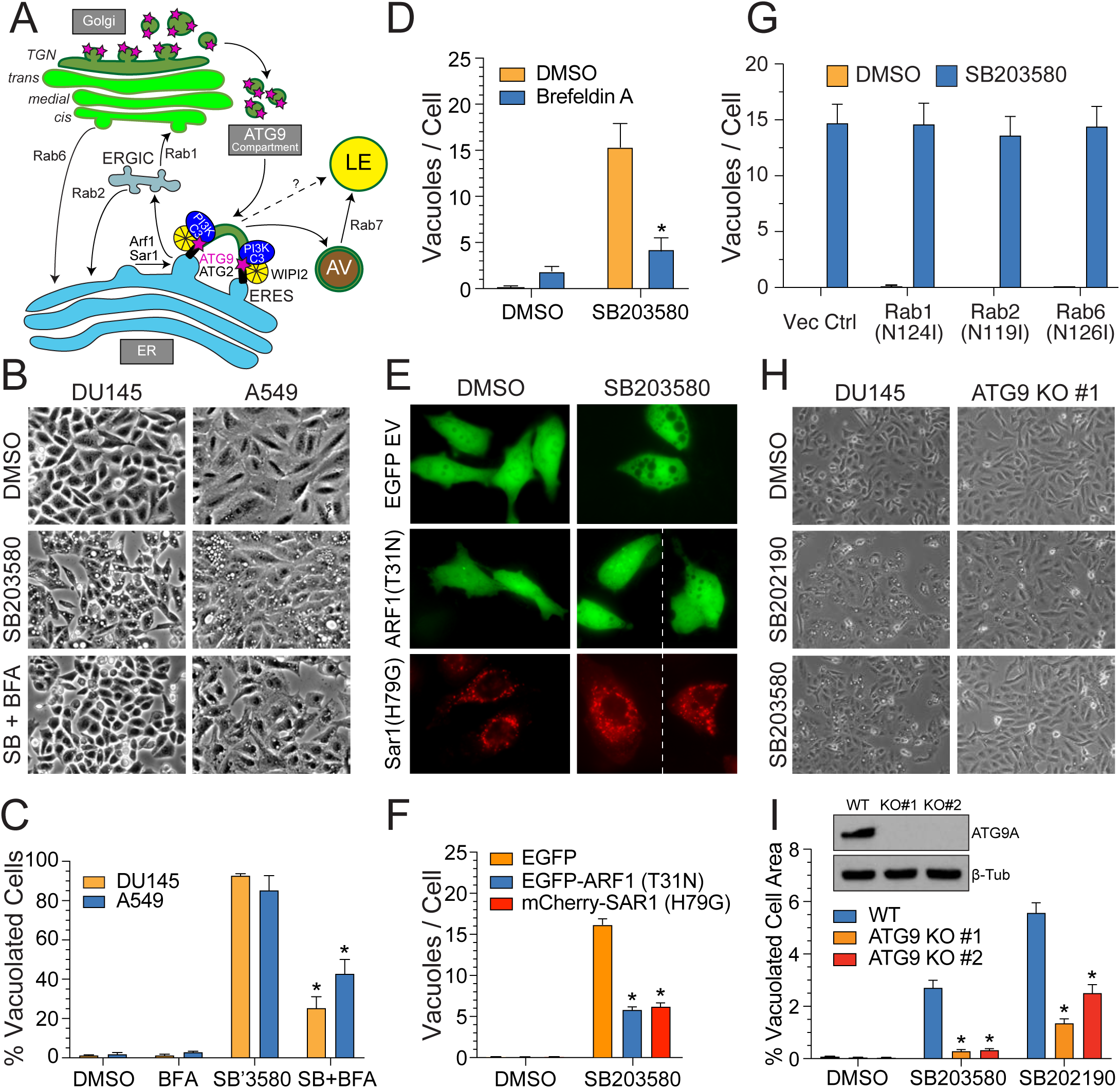
Selective disruption of ERES and ATG9A inhibits SB203580-induced vacuolation. *(A)* Cartoon depicting the roles of various ARF and RAB GTPases in ER-Golgi trafficking, as well as the importance of ERES and ATG9A in mediating the formation of PAS. *(B-D)* DU145 and A549 cells were treated with SB203580 (50 µM) ± BFA (10 µg/mL) and assessed for vacuolation by phase-contrast microscopy. *(E,F)* DU145 cells were transfected with either EGFP (vector control), dominant-negative Arf1(T31N)-EGFP, constitutively active mCherry-Sar1(H79G), or *(G)* dominant-negative mCherry-Rab1(N124I), Rab2(N119I), or Rab6(N126I). The cells were then treated with DMSO or SB203580 (50 µM) and assessed for vacuolation by fluorescence microscopy. *(H,I)* CRISPR-Cas9 was utilized to delete *ATG9A* from DU145 cells, and two separate clones were then exposed to SB203580 (50 µM) or SB202190 (50 µM) and assessed for vacuolation by phase-contrast microscopy.

The small GTPases, ARF1 and SAR1, mediate the budding and transport of vesicular membranes between the ER and Golgi compartments, and the fungal metabolite Brefeldin A (BFA) directly inhibits ARF1 by locking it into its GDP-bound form, resulting in the collapse of the Golgi apparatus and the disruption of ERES (9,40). Therefore, we pretreated cells with BFA and observed a significant decrease in SB203580-induced vacuolation, both in the number of vacuolated cells and in the number of vacuoles per cell, in autophagy-deficient (DU145) and autophagy-competent (A549) cells (Fig. 4B-D). Dominant-negative GDP-locked ARF1(T31N) and constitutively-active GTP-locked SAR1(H79G) also disrupt ERES (9), and similar to BFA treatment, overexpression of these mutants significantly inhibited the formation of SB203580-induced vacuoles (Fig. 4E,F). Conversely, overexpression of Rab1, Rab2, or Rab6 dominant-negative constructs, which inhibit various aspects of anterograde and retrograde trafficking between ER and Golgi (41), failed to suppress vacuolation (Fig. 4G).

The essential transmembrane autophagy protein, ATG9A, is incorporated into vesicles that emanate from the TGN (13,42), and these vesicles, dubbed the “ATG9A compartment”, are subsequently recruited to PAS at ERES, where they serve as the initial membrane source for the growth of phagophores (Fig. 4A) (15). p38 MAPKs are also thought to regulate ATG9A function through phospho-dependent sequestration of p38IP, an ATG9A interactor that influences its trafficking between the TGN, endosomes, and PAS (17). Mechanistically, ATG9A is a lipid scramblase, which, in conjunction with ATG2, mediates the transfer and interleaflet redistribution of phospholipids from ERES to expanding phagophores (Fig. 4A) (14,43,44). Since the collapse of Golgi and blockage of ERES suppressed SB203580-induced vacuolation, we questioned whether ATG9A-dependent phagophore membranes might contribute to vacuole formation (Fig. 4A). We therefore deleted ATG9A from autophagy-deficient DU145 cells using CRISPR-Cas9 and found that, indeed, loss of ATG9A substantially inhibited vacuolation (Fig. 4H,I).

Collectively, the data strongly suggested, at a minimum, that the ER/Golgi compartments represented significant membrane sources for the growth of vacuoles following SB203580 treatment and did so by way of ERES and ATG9A. In autophagy-competent cells, this likely occurs through the fusion of autophagosomes with LELs (Fig. 2F-I). However, it is worth noting that ATG9A colocalizes with Rab7 on endosomal membranes (45), and ER-endosome “contact sites” are known to facilitate the direct transfer of certain lipids and proteins from ER to late endosomes (38,46–48). Besides the juxtaposition of ER membranes to the expanding SB203580-induced vacuoles (fig. S4A), we also observed an apparent transport of *cis*-Golgi (GRASP65 and GOS28), but not ERGIC (ERGIC53), *medial*-Golgi (GRASP55), or *trans*-Golgi (TGN38) proteins to the enlarged LELs (fig. S4A-C). Thus, while more formal proof is beyond the scope of the current study, we cannot rule out the possibility that the ATG2/ATG9A complex and ER-endosome contact sites may mediate the direct transfer of lipids and proteins from remodeled ERES to the growing vacuole (Fig. 4A).

### SB203580 disrupts the osmotic balance and function of LELs

p38 MAPK inhibitors were initially reported to induce autophagy in large part due to an apparent increase in LC3B lipidation (18), which we too observed in autophagy-competent cells, including HT-29, HCT116, and A549 cells (Fig. 5A, lanes 1 and 2). However, given that SB203580 failed to stimulate LLPD in these same cell lines (fig. S2C), we speculated that the observed increase in lipidated LC3B (LC3-II) might have resulted instead from a decrease in autophagic flux, similar to that observed following alkalinization of LELs with NH_4_Cl (Fig. 5A, lanes 2 and 4). Moreover, treatment of cells with SB203580 and SB202190 caused a decrease in the pre-pro processing of cathepsin D (CTSD) and a block in the degradation of p62 (Fig. 5B). Since cathepsin activity is pH-dependent, we assumed that SB203580 treatment resulted in LEL alkalinization; instead, we discovered that SB203580 stimulated the formation of numerous highly acidic vesicles that readily stained with both LysoTracker^TM^ Green and Red dyes (Fig. 5C,D). These acidic vesicles were not clearly visible by phase-contrast imaging but did surround the much larger translucent and more weakly stained vacuoles (Fig. 5C). This led us to speculate that fusion of the smaller acidic vacuoles might have given rise to the much larger, less acidic vacuoles. Therefore, we pretreated cells with Bafilomycin A (Baf A), a potent inhibitor of the V-ATPase that alkalinizes LELs and inhibits their fusion with one another and autophagosomes (49–51). As predicted, Baf A profoundly suppressed both SB203580-induced cellular acidification and vacuolation, as did direct alkalinization with chloroquine (Fig. 5D,E; fig. S5A).

**Figure 5.**
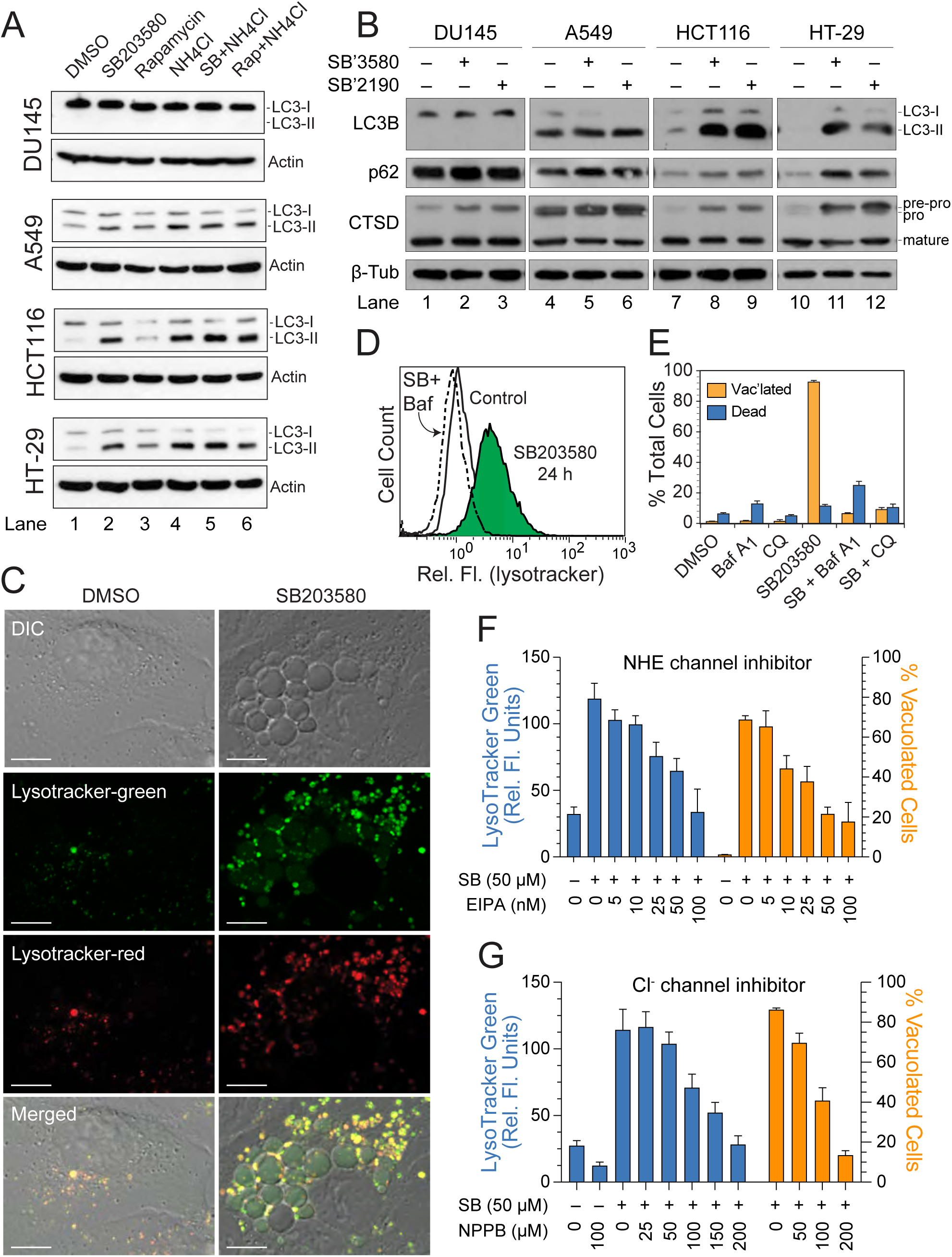
SB203580 induces LEL swelling and defects in cathepsin processing and protein degradation. *(A,B)* DU145, A549, HCT116, and HT-29 cells were exposed to SB203580 (50 µM), SB202190 (50 µM), or rapamycin (200 nM) for 24 h, plus or minus NH_4_Cl, and then western blotted for LC3B, p62, CTSD, or β-tubulin. *(C,D)* DU145 cells were exposed to SB203580 (50 µM) for 24 h, labeled with LysoTracker^TM^ Green and/or Red, and evaluated by confocal microscopy or flow cytometry. *(E-G)* DU145 cells were treated with SB203580 (50 µM) for 24 h, in the presence or absence of a V-ATPase inhibitor (Bafilomycin A1, 125 nM), an alkalinizing lysosomatropic agent (chloroquine, 10 µM), a Na+-H+ exchanger inhibitor (ethylisopropyl amiloride, *i.e.* EIPA, 5-100 nM), or a chloride channel inhibitor (5-nitro-2-(3-phenylpropyl-amino)benzoic acid, *i.e.* NPPB, 25-200 µM). The cells were then assayed for acidic compartments by flow cytometry (LysoTracker^TM^ Green) and evaluated for vacuole formation by phase-contrast microscopy.

While acidification of vesicles is required for membrane fusion, we noted that the larger translucent vacuoles were not only less acidified but also exhibited signs of increased swelling, perhaps due to increased turgor pressure (Fig. 5C; Fig. 2E; fig. S2B,D). Indeed, following washout of SB203580, LysoTracker^TM^ Green staining of cells rapidly returned to normal (fig. S5B,C); and the vacuoles underwent a collapse, with some exhibiting apparent signs of pinching off (fig. S5C, *images iii-vi*). This led us to question whether an initial acidification of smaller vesicles facilitated membrane fusion but eventually led to an influx of water that resulted in severe vacuole (*i.e.* LEL) swelling. Consistent with this interpretation, incubation of cells in hypertonic medium containing sorbitol dramatically reduced SB203580-induced vacuolation (fig. S5D); and addition of Baf A to already vacuolated cells also resulted in vacuole resolution (fig. S5E). Mechanistically, several ion transporters, including Na^+^/H^+^ exchangers (NHEs) and Cl^-^/H^+^ exchangers (CLCs), exchange protons for sodium or chloride ions, resulting in an influx of salt into the lumens of LELs (fig. S5F). With this in mind, we pretreated cells with the NHE inhibitors, EIPA or Zonaporide, or the chloride channel inhibitors, NPPB or DIDS, and observed concentration-dependent decreases in LysoTracker^TM^ Green staining and/or vacuole formation (Fig. 5F,G; fig. S5G,H). These effects appeared to be specific, since inhibitors of the CFTR chloride channel or Na^+^/Ca^2+^ exchangers had no significant effects (fig. S5I,J). We were, of course, aware that EIPA is an excellent inhibitor of macropinocytosis, but as already noted, SB203580 potently inhibited acute uptake of the fluid-phase markers, Dextran-FITC and BSA-FITC (fig. S3D,E). Thus, the profound effects of both Baf A and EIPA on SB203580-induced vacuolation were most likely attributable to an inhibition in the proton-driven influx of NaCl and water into the lumens of LELs. Altogether, the data to this point suggested that SB203580 stimulated Rab7-dependent fusion of ER/Golgi-derived membranes (including autophagosomes) with LELs; however, severe swelling of these LELs hindered both cathepsin processing/activity and degradation of any delivered cargo due to increased volume and/or alkalinization of the enlarged compartment.

### SB203580 and SB202190 stimulate vacuolation through off-target inhibition of PIKfyve

During our efforts to uncover the mechanisms responsible for SB203580-induced vacuolation, we noted striking phenotypic similarities between the vacuoles induced by pyridinyl imidazole p38 MAPK inhibitors and the more recently developed inhibitors of the lipid kinase PIKfyve, which catalyzes the conversion of PI to PI(5)P and PI(3)P to PI(3,5)P2 (52–56). PI(3,5)P2, in particular, is known to positively regulate the activities of at least two classes of ion channels: ‘transient receptor potential’ (TRP) channels, including TRPML1 (also known as MCOLN1), and ‘two pore channels’ (TPCs) (57–63). Thus, PIKfyve inhibition and the subsequent decrease in PI(3,5)P2 is reportedly associated with dysregulation of calcium and sodium efflux, LEL membrane potential, and ultimately LEL intralumenal invagination, swelling, and fission/fusion dynamics (56,64–66). PIKfyve inhibitors, YM201636 and apilimod, stimulated the formation of vacuoles that appeared largely indistinguishable from those induced by SB203580 and SB202190 (Fig. 6A; Fig. 2E; fig. S2B,D; fig. S6A). This led us to question whether SB203580 and SB202190 might induce vacuolation by directly inhibiting PIKfyve, and indeed, while less potent than YM201636, both SB203580 and SB202190 inhibited recombinant PIKfyve kinase activity *in vitro* with IC_50_ values of 433 ± 42.9 nM and 355 ± 36.2 nM, respectively (Fig. 6B). SB203580 and SB202190 likewise reduced the levels of PI(3,5)P2 in treated cells (Fig. S6B), as determined by immunofluorescence staining (22). SB203580 and SB202190 also induced a block in cathepsin processing and autophagic flux that mirrored that of other PIKfyve inhibitors (Fig. 6C). Conversely, the structurally unrelated p38 MAPK inhibitor, BIRB-796, failed to inhibit PIKfyve, induce vacuolation, suppress cathepsin processing, or inhibit autophagic flux, despite inhibiting the p38 MAPK pathway (Fig. 6A-C; Fig. S6C). Thus, collectively, the data suggested that pyridinyl imidazole p38 MAPK inhibitors induced vacuolation through “off-target” inhibition of PIKfyve.

**Figure 6.**
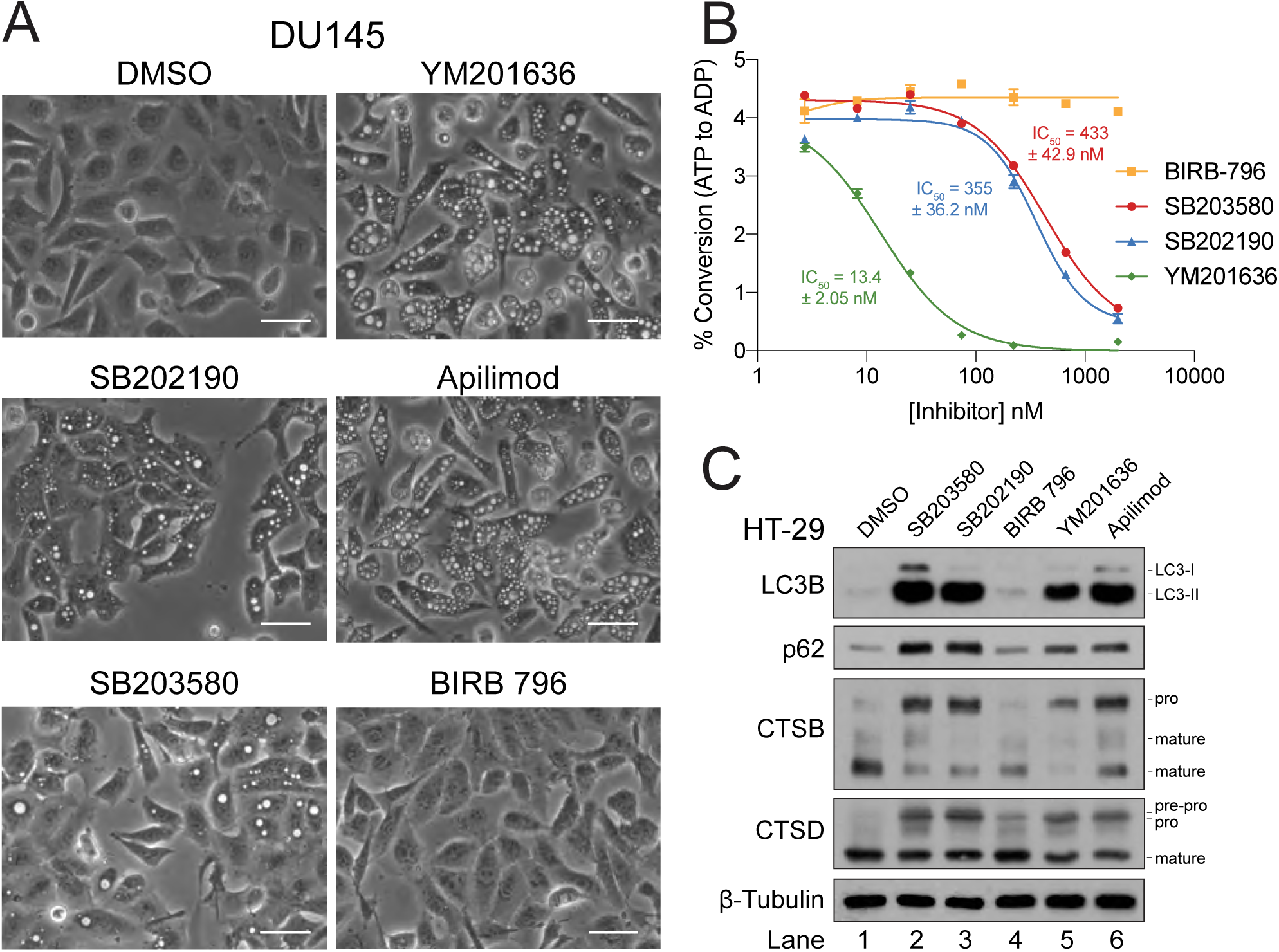
Pyridinyl imidazole p38 MAPK inhibitors are direct inhibitors of the lipid kinase PIKfyve. *(A)* DU145 cells were treated with SB202190 (50 µM), SB203580 (50 µM), YM201636 (1 µM), apilimod (50 nM), or BIRB-796 (50 µM) for 24 h and then evaluated for vacuolation by phase-contrast microscopy. *(B)* Recombinant PIKfyve was incubated with PI(3)P, phosphatidylserine, and ATP, in the presence of increasing concentrations of SB203580, SB202190, BIRB-796, or YM201636, and assayed for ATP hydrolysis using ADP Glo luminescence reagent, as described in the methods. *(C)* Autophagy-proficient HT-29 cells were treated with SB203580 (50 µM), SB202190 (50 µM), BIRB-796 (50 µM), YM201636 (1 µM), or apilimod (50 nM) for 24 h and then immunoblotted for LC3, p62, CTSB, CTSD, and β-tubulin.

### p38 MAPKs suppress vacuolation triggered by PIKfyve inhibition by promoting LEL fission

The fact that SB203580, but not BIRB-796, could inhibit PIKfyve and induce vacuolation ostensibly ruled out a role for p38 MAPK inhibition in vacuolation. However, we had already demonstrated that a constitutively-active p38α, containing the T106M gate-keeper mutant, could significantly rescue cells from SB203580-induced vacuolation (Fig. 1F). This led us to question whether SB203580 and SB202190 might stimulate vacuolation *via* the simultaneous inhibition of both PIKfyve and p38 MAPKs (Fig. 7A). To evaluate whether p38 MAPKs might regulate vacuolation epistatic to PIKfyve, we generated p38α/β double knockout (p38 DKO) DU145 cell lines using CRISPR/Cas9 (Fig. 7B) and treated both wild-type (WT) and p38 DKO cells with SB203580, SB202190, YM201636, or apilimod (Fig. 7C,D). Remarkably, vacuolation was dramatically more severe in p38 DKO cells, following all treatments, with the cytoplasm of some cells being almost entirely occupied by a single very large vacuole (Fig. 7C,D). Acute pharmacological inhibition of p38 MAPK activities with BIRB-796 in wild-type DU145 cells likewise exacerbated YM201636 and apilimod-induced vacuolation (fig. S7A-C), further indicating that combined inhibition of PIKfyve and p38 MAPKs induced more extensive vacuolation than PIKfyve inhibition alone.

**Figure 7.**
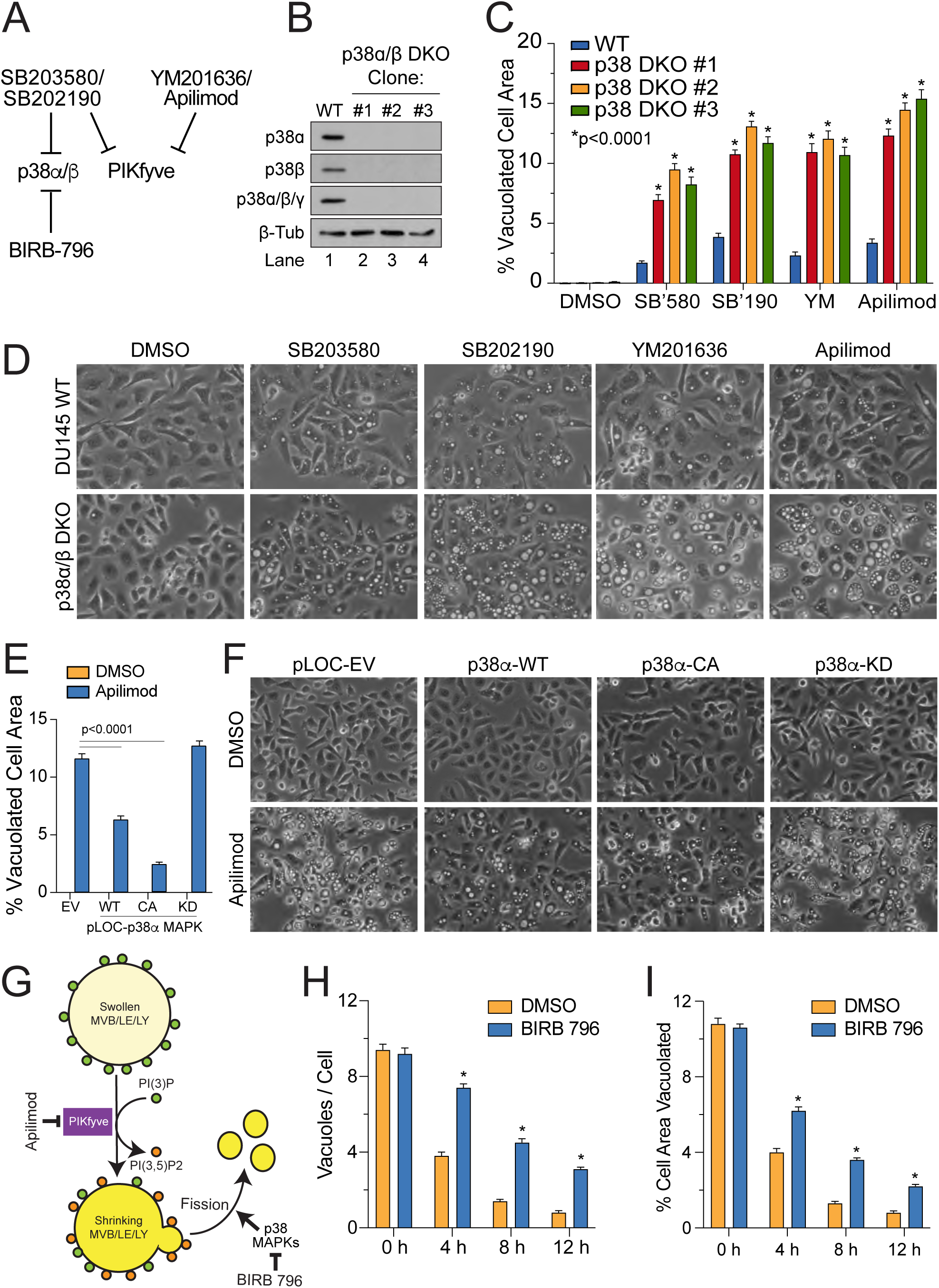
Loss or inhibition of p38 MAPKs sensitizes cells to vacuolation induced by PIKfyve inhibitors. *(A)* Model to explain how SB203580 and SB202190 induce vacuolation through combined inhibition of p38 MAPKs and PIKfyve. *(B)* p38α and p38β were deleted from DU145 cells using CRISPR-Cas9; and *(C,D)* three p38 DKO clones were exposed to SB203580 (50 µM), SB202190 (50 µM), YM201636 (500 nM), or apilimod (20 nM) and evaluated for vacuolation by phase-contrast microscopy using ImageJ analysis software. *(E,F)* p38 DKO cells were stably reconstituted with a vector control, or wild-type (WT), constitutively-active (CA), or kinase dead (KD) versions of p38α, exposed to apilimod (20 nM), and evaluated for vacuolation by phase-contrast microscopy using ImageJ analysis software. *(G)* Model for epistatic relationship between PIKfyve and p38 MAPKs. *(H,I)* Wild-type DU145 cells were treated with apilimod (50 nM) for 24 h, after which apilimod was removed and BIRB-796 (50 µM) was added to selectively inhibit p38 MAPKs during the “washout” phase. Vacuolation was assessed by phase-contrast microscopy over time (0-12 h) using ImageJ software to determine both the number of vacuoles per cell and percent cell area vacuolated (see also fig. S7D). Inhibition of p38 MAPKs delays the resolution of vacuoles, indicating a role for p38 MAPKs in promoting LEL fission.

Notably, the fact that p38 DKO cells (compared to wild-type cells) still exhibited increased vacuolation following treatment with SB203580 and SB202190, even though no p38 MAPKs were present, suggested that complete loss of p38 MAPKs may have resulted in better or more sustained inhibition than could be pharmacologically achieved in wild-type cells (Fig. 7C,D). However, we could not rule out the possibility that p38 MAPK proteins played a structural/scaffolding role in suppressing vacuolation as opposed to their activities alone. Therefore, we reintroduced wild-type (WT), constitutively-active (CA), or kinase dead (KD) p38α MAPKs into DKO cells and exposed them to apilimod (Fig. 7E,F). As anticipated, expression of p38α-WT and particularly p38α-CA suppressed apilimod-induced vacuolation; however, expression of p38α-KD had no apparent effect (Fig. 7E,F), reaffirming that p38 MAPK activity *per se* was required to suppress vacuolation induced by PIKfyve inhibition.

Collectively, these data led us to speculate that p38 MAPKs might play a role in *promoting* LEL fission (Fig. 7G). If so, we reasoned that the rate at which vacuoles resolve, following the removal of PIKfyve inhibitor, should be reduced when p38 MAPKs are selectively inhibited (Fig. 7G). We therefore treated DU145 cells with apilimod to induce vacuolation; and cells were then washed, incubated with either DMSO or BIRB-796, and evaluated over 12 h for vacuolation. Following “washout”, BIRB-796 significantly delayed the resolution of vacuoles, indicating that p38 MAPKs did indeed play a significant role in LEL fission (Fig. 7H,I; fig. S7D). Thus, p38 MAPKs act epistatic to PIKfyve with regard to LEL fission-fusion dynamics; and pyridinyl imidazole p38 MAPK inhibitors are effective at inducing to cytoplasmic vacuolation due to their combined “on-target” inhibition p38 MAPKs and “off-target” inhibition of PIKfyve.

## DISCUSSION

In this study, we observed cytoplasmic vacuolation in cells exposed to pyridinyl imidazole p38 MAPK inhibitors, SB203580 and SB202190; and we confirmed that p38α was an important target using a drug-resistant “gatekeeper” mutant that suppressed vacuolation (Fig. 1; fig. S1). However, the structurally unrelated p38 MAPK inhibitor, BIRB-796, failed to induce vacuolation, despite the inhibition of p38 MAPKs (Fig. 6), creating a conundrum that hampered progress for some time. We surmised that p38 MAPK inhibition alone was required but insufficient to induce vacuolation and that SB203580 must have another target whose *simultaneous* inhibition was required for vacuolation. Thus, we were thrust into utilizing a modified phenotypic drug discovery approach to identify the “off-target” of SB203580 responsible for vacuolation in the context of p38 MAPK inhibition (67).

During our studies, published reports suggested that vacuolation induced by SB202190 resulted from an increase in autophagic cell death (18,19). However, while PIK3C3 was essential for SB203580-induced vacuolation – and autophagosomes resulting from basal autophagy could contribute to vacuole enlargement – SB203580 itself did not induce autophagy *per se*, at least as defined by an increase in the degradation of cellular cargo (Fig. 2; fig. S2). In fact, SB203580 suppressed autophagic flux, resulting in an accumulation of LC3-II and p62 in autophagy-competent cells (Fig. 5). Since vacuolation might theoretically result from any number of defects in endomembrane trafficking, we sought to characterize the nature of the vacuoles and identified them as severely swollen LELs (Fig. 3). Growth of these vacuoles depended in part on Rab7, as well as ERES activity and ATG9A-dependent flow of membrane from the ER/Golgi compartments (Figs. 3 and 4). However, enlargement of these vacuoles was driven by an osmotic imbalance in LELs, resulting from an initial hyper-acidification, followed by a subsequent influx of salt and water (Fig. 5; fig. S5).

By this stage of the project, PIKfyve inhibitors were reported in the literature to induce the formation of cytoplasmic vacuoles that were remarkably similar to those we observed with SB203580. We subsequently determined that SB203580 and SB202190 could, in fact, inhibit PIKfyve *in vitro*, albeit with lower potency, and reduced the production of PI(3,5)P2 in cells (Fig. 6B; Fig. S6B). Finally, using CRISPR/Cas9-derived p38 DKO cells and a series of pharmacological experiments, involving selective PIKfyve inhibitors and BIRB-796, we confirmed that p38 MAPKs played a critically important role in mediating LEL fission (Fig. 7; fig. S7). Thus, pyridinyl imidazole p38 MAPK inhibitors induced cytoplasmic vacuolation through a combination of PIKfyve and p38 MAPK inhibition, resulting in profound LEL swelling and a reduction in LEL fission. Our study underscores the inherent challenges of phenotypic drug discovery, particularly in instances where a compound possesses activities on different targets at distinct steps in the same pathway. However, our characterization of SB203580 has revealed an epistatic relationship between PIKfyve and p38 MAPKs that would likely have been difficult to uncover otherwise. Indeed, loss of p38 MAPKs (or inhibition of them with BIRB-796) was insufficient to induce vacuolation on its own; and complete inhibition of PIKfyve activity results in profound vacuolation that can mask the effects of p38 MAPK inhibition. It is therefore only in the context of weaker PIKfyve inhibition that p38 MAPK inhibition reveals its true impact on cytoplasmic vacuolation.

Importantly, there are at least two practical implications from our study: First, considering the recent interest in the clinical use of PIKfyve inhibitors, particularly for the treatment of various cancers (55,56,68), and given that complete intratumoral inhibition of PIKfyve or p38 MAPK activities is unlikely *in vivo*, combined use of PIKfyve and p38 MAPK inhibitors may prove therapeutically useful in the future. This is particularly true, since p38 MAPK pathways are reportedly activated by PIKfyve inhibitors in cancer cells (69). Second, although SB203580 and SB202190 are themselves unlikely to be utilized clinically, it is worth emphasizing from a basic science perspective that both compounds have frequently been employed to evaluate the roles of p38 MAPKs in various endomembrane trafficking processes including endocytosis and autophagy. Many of these processes are likely to be impacted by unexpected off-target inhibition of PIKfyve, so care should be exercised when using these compounds and results from previous studies may warrant reevaluation with this fact in mind.

Finally, while our phenotypic drug discovery approach to SB203580-induced vacuolation identified PIKfyve as a critical target (and helped demonstrate its epistatic relationship to p38 MAPKs), the specific downstream target(s) of p38 MAPKs that regulate LEL fission remain unknown and are the focus on ongoing work. PIKfyve inhibition clearly disrupts normal osmotic balance in LELs and induces swelling; and a decrease in PI(3,5)P2 levels is known to reduce the activities of various ion channels, including TRPML1, TPCs, and V-ATPases in mammals and/or yeast (57–63). Thus, it is possible that p38 MAPKs positively regulate the activities of one or more of these channels (or others), some of which possess putative phosphorylation sites, and in turn promote LEL fission by reducing swelling. Other potential targets clearly exist, including those proteins that constitute the putative lysosome-specific fission machinery such as WIPI, SNX, and dynamin proteins (70–72). Given the relatively large number of conceivable targets, a phospho-proteomic analysis of wild-type and p38 MAPK-deficient (or acutely inhibited) cells, analyzed following exposure to PIKfyve inhibitors and during washout of the drug, may offer the best opportunity to identify those targets of p38 MAPKs that are critical for LEL fission.

## MATERIALS AND METHODS

### Reagents

SB203580, SB202190, SP600125, PD98059, LY294002, Bafilomycin A1, and rapamycin were obtained from Alexis Corp. (San Diego, CA) or Tocris Cookson (Ellisville, MO). All other chemicals were of analytical grade and purchased from Sigma-Aldrich or EMD Biosciences (Gibbstown, NJ). Antibodies to ATG5 (#2630), ATG12 (#4180), ATG16L1 (#8089), p62 (#5114), LCB (#2775), Beclin 1 (#3738), ATG9A (#13509), CTSD (#2284), CTSB (#31718), β-actin (#4970), p38α (#9217), p38β (#2339), phospho-p38α (pThr-180/pThr-182; #9216), phospho-MK2 (pThr-222; #3316) and phospho-hsp27 (pSer-82; #2406) were purchased from Cell Signaling Technology (Danvers, MA). Anti-PI(3,5)P2 (Z-P035) was obtained from Echelon Biosciences (Salt Lake City, UT). Anti-tubulin (RE11250C100) was obtained from BioVendor (Candler, NC). For immunofluorescence experiments, LAMP1 (H4A3) was purchased from Developmental Studies Hybridoma Bank at the University of Iowa (Iowa City, IA). Alexa Fluor 488- and HRP-conjugated secondary antibodies to rabbit and mouse IgG were obtained from Molecular Probes^®^ Invitrogen (Carlsbad, CA), DakoCytomation, and Sigma-Aldrich (St. Louis, MO).

### Plasmid constructs

A cDNA encoding p38α was subcloned into pcDNA6 with an N-terminal myc tag (Invitrogen, Carlsbad, CA), and a kinase-dead (D168A) mutant was subsequently generated by site-directed mutagenesis. A constitutively-active mutant of p38α (D176A/F327S; obtained from Prof. David Engelberg, Hebrew University, Jerusalem) (73) was subcloned into the *Xho*I-*Bam*HI sites of the pEGFP-C1 vector (Clontech, Mountain View, CA), and the corresponding SB203580-resistant mutant (T106M) was generated by site-directed mutagenesis. An shRNA to human *beclin 1* (5’-1206-GATTGAAGACACAGGAGGC-1225-3’) was cloned into the *Bgl*II/*Xho*I sites of pSuper.retro.puro, transfected in DU145 cells and selected for stable cells using puromycin. A plasmid encoding EGFP-LC3 was provided by Prof. A. Thorburn (University of Colorado Health Sciences Center, Denver, CO) (29).

Labeling of early endosomes was performed by transfecting cells with EGFP-Rab5 or EGFP-EEA1 constructs (kindly provided by Prof. M. Seabra, Imperial College, UK, and Dr. S. Corvera, University of Massachusetts Medical School, Worcester, MA) (74,75). Dominant-negative Rab5 (N133I) and the p38α-specific phosphomimetic EEA1 mutant (T1392E) were subsequently generated by site-directed mutagenesis. Human GDIα was obtained by RT-PCR (forward: 5’-CTCGAGCTATGGACGAGGAATACGACGTGA-3’; reverse: 5’-GGATCCTCACTGGTCAGCTTCCCCAAA-3’) and cloned into pmCherry-C1, and its phosphomimetic mutant (S121E) was later cloned by site-directed mutagenesis. Labeling of LEs was performed by transfecting cells with EGFP-Rab7 or EGFP-Rab9 (provided by Dr. B. V. Deurs, University of Copenhagen, Denmark, and Prof. S. Pfeffer, Stanford University, Palo Alto, CA) (76,77). Dominant-negative Rab7 (T22N) and Rab9 (S21N) mutants were then generated by site-directed mutagenesis. For certain co-labeling experiments, the Rab7 and Rab9 markers were also subcloned into pmCherry-C1 in order to generate Cherry fusion proteins. Finally, lysosomes were labeled using LAMP1-mGFP or EGFP-CD63 constructs (provided by Dr. E. Dell’Angelica, UCLA, CA, and Dr. P. Luzio, University of Cambridge, UK) (78,79).

In order to visualize the generation of PI3P, the PX domain of p40Phox (aa 1-149) was cloned by RT-PCR (forward: 5’-GGTACCGAATGGCTGTGGCCCAGCAGC-3’; reverse: 5’-CTCGAGGCGGATGGCCTGGGGCACC-3’) into the *Kpn*I/*Xho*I sites of pcDNA3, in frame with a C-terminal EGFP tag (*Xho*I/*Xba*I) (28). The PX mutant R57Q, which does not bind to PI3P, was subsequently generated by site-directed mutagenesis. For labeling ER membranes, a cDNA encoding the C-terminus (aa 100-134) of human cyt. b5, was obtained by RT-PCR (forward: 5’-CTCGAGGAATGATCACTACTATTGATTCTAG-3’; reverse: 5’-GGATCCTCAGTCCTCTGCCATGTATAGG-3’) and cloned into the *Xho*I-*Bam*HI sites of pEGFP-C1. ER-Golgi intermediate carriers (ERGICs) were labeled using ERGIC53-GFP (provided by Prof. Hans-Peter Hauri, University of Basel, Switzerland) (80); and *cis*, *medial* or *trans*-Golgi membranes were labeled with GRASP65-EGFP, EGFP-GOS28, GRASP55-EGFP, and TGN38-EYFP (provided by Prof. Francis Barr, University of Liverpool, Liverpool, UK; and Dr. Walther Mothes, Yale University, New Haven, CT) (81,82).

The cDNA of human ARF1 was obtained by RT-PCR (forward: 5’-CTCGAGATGG GGAACATCTTCGCCAACC-3’; reverse: 5’-GGATCCGGCTTCTGGTTCCGGAGCTGA-3’) and cloned into the *Xho*I-*Bam*HI sites of pEGFP-N1, whereas human SAR1 was cloned (forward: 5’-CTCGAGCTATGTCTTTCATCTTTGAGTGGA-3’; reverse: 5’-GGATCCTCAGTCAATATACTGGGAGAGCC-3’) into the *Xho*I-*Bam*HI sites of pmCherry-C1. Dominant-negative Arf1(T31N) and constitutively-active Sar1(H79G) mutants were then generated by site-directed mutagenesis. Rab1, Rab2, and Rab6 cDNAs (kind gift of Dr. Terence Hébert, McGill University, Canada) (83) were subcloned into the *Xho*I-*Bam*HI sites of pmCherry-C1, and the corresponding dominant-negative mutants were generated by site-directed mutagenesis.

The pLOC lentiviral expression vector was obtained from the University of Texas MD Anderson Cancer Center (UTMDACC) Functional Genomics Core (FGC). Full-length ATG5 was cloned into BamHI/NheI sites of untagged pLOC and pLOC-HA. Wild-type, kinase dead (D168A) and constitutively-active (D176A/F327S) p38α mutants were cloned into the same sites.

### Cell culture and transfections

All cancer cell lines and MEFs were grown in RPMI-1640 or DMEM, supplemented with 5% fetal bovine serum (Atlanta Biologicals, Norcross, GA), 5% Fetalplex (Gemini Bio-products, West Sacramento, CA), 1% penicillin-streptomycin (100 units/mL) and 2 mM Glutamine. Cells were maintained at 37°C in humidified air containing 5% CO_2_ and were routinely passaged every 3 days. For all transient transfections, with the exception of EGFP-LC3, cells were transfected with 1 µg/mL plasmid DNA using Fugene HD transfection reagent (Roche Diagnostics, Indianapolis, IN).

### Lentiviral transduction and stable cell line generation

HEK293T cells were co-transfected with pLOC along with psPAX2 and pHCMV-G lentivirus packaging plasmids. psPAX2 was a gift from Dr. Didier Trono (Addgene plasmid #12260). Approximately 48 h following transfection, the medium was collected and 6 μL of sterile hexadimethrine bromide (5 μg/μL; Sigma-Aldrich #H9268) was added. The collected medium was then filtered through a 0.45 μm PVDF syringe filter and incubated with target cells overnight. Transduction efficiency was evaluated by GFP expression and/or immunoblotting.

### Cell proliferation, endo-lysosomal volume, and apoptosis Assays

To measure cell proliferation, cells were labeled with CFDA-SE (10 µM; Molecular Probes) for 15 min. Excess CFDA-SE was removed by washing the cells three times with culture media at 25°C. Labeled cells were then plated in 12-well dishes, treated with SB203580 for several days, and analyzed by flow cytometry (Cytomics FC 500, Beckman-Coulter) for the residual fluorescence in dividing cells. To measure endo-lysosomal volume, cells were treated for 24-48 h with DMSO (vehicle control) or SB203580, trypsinized, labeled with LysoTracker^TM^ Green DND-26 (50 nM; Molecular Probes) for 60 min at 37°C, and analyzed by flow cytometry. Apoptosis was assessed by Annexin V/propidium iodide (PI) staining, as previously described (84).

### Vacuolation assays and transmission electron microscopy

To determine the effects of various kinase inhibitors on cytoplasmic vacuolation, DU145 prostate cancer cells were treated with 50 µM of SB203580 or SB202190 (p38), SP600125 (JNKs), PD98059 (ERKs), LY294002 (PI3 kinases), or rapamycin (mTOR). In addition, cells were treated with SB203580 (50 µM) for 24 h in the presence or absence of Baf A_1_ (125 nM) or CQ (10 µM). The cells were then examined by phase-contrast light microscopy, and the percentage of vacuolated cells was determined after counting at least 200 cells from three random fields. Each experiment was performed at least three times, and each data point represents mean ± SEM. In experiments performed later in the project (e.g. Figs. 6 and 7), cells (>1,000) were analyzed using ImageJ software and macro developed in-house to determine both vacuole counts and total vacuolated area per cell. For transmission electron microscopy, cells were cultured in 12-well dishes, treated with SB203580 for 24 h, fixed with 1.5% buffered glutaraldehyde and 1% formaldehyde, post-fixed in 2% CaCo-buffered osmium tetroxide-0.8% potassium ferricyanide, embedded in EPON™ resin, sectioned, and analyzed using a Philips EM 208 transmission electron microscope.

### Long-lived protein degradation (LLPD) assay

Cells were labeled with 0.2 μCi/mL L-valine [^14^C(U)] (MC-277, Moravek Biochemicals, Brea, CA) overnight. Media was then collected and cells were washed 3 times with Hank’s buffered saline solution (HBSS). Cells were then chased with fresh media containing 10 mM L-valine for 1 h. Control (t = 0) cells were immediately harvested and precipitated with 10% Trichloroacetic Acid (TCA). Precipitates were spun down and the soluble fraction was removed and collected while the insoluble fraction was dissolved in 1 mL 0.2 M NaOH. The rest of the treated cells were washed twice with HBSS and then incubated with media containing 10 mM L-valine and one of the following treatments: dimethyl sulfoxide (DMSO), 3-methyladenine (3-MA) (10 mM), HBSS + DMSO or HBSS + 3-MA (10 mM). At each time point 50 μL of media from each well was collected and TCA precipitated with 50 μL of 20% TCA. A portion of each fraction (100 μL) was added to 5mL of scintillation cocktail (Biocount cocktail, RPI Corp #111182) and assayed using a scintillation counter. The *total radioactivity (T)* was calculated by adding the measured counts per minute (cpms) of the soluble and insoluble fractions at t = 0 and multiplying by 10 (1:10 dilution factor). The measured cpms for the treated cells at each time point were multiplied by 40 (1:40 dilution factor) and divided by the *total radioactivity (T)* to give a % long-lived protein degradation. To isolate degradation specifically related to autophagy from other forms of protein turnover, we subtracted the cpms from 3-MA treated cells from the cpms from DMSO treated cells. This results in the % of 3-MA sensitive protein degradation.

### PIKfyve Inhibition Assay

recombinant PIKfyve was incubated with inhibitors (SB203580, SB202190, YM201636, or BIRB-796) in kinase assay buffer (50 mM HEPES pH 7.5, 50 mM NaCl, 20 mM MgCl_2_, 0.5 mM EGTA, 0.04% Triton X-100, 25 µg/mL BSA, 0.05 mM DTT) for 30 min at room temperature. Kinase reactions were initiated by adding 5 µL of a mixture of PI(3)P, phosphatidylserine (PS), and ATP. After 2 h, 5 µL of ADP Glo Reagent (Promega), supplemented with 0.01% Triton X-100, was added and incubated for an additional 40 min. Finally, Kinase Detection Reagent (10 µL) was added, incubated for an additional 30 min, and each sample assayed for luminescence. Final assay concentrations: PIKfyve (1 ng/µL), PI(3)P (50 µM), PS (400 µM), ATP (50 µM), inhibitor (0-2,000 nM), and DMSO (0.02%).

### Immunofluorescence and western blotting

Cells grown on coverslips were fixed in 4% paraformaldehyde for 15 min, followed by a 5 min fixation in ice-cold methanol and an overnight block in PBS containing 10% goat serum. The fixed cells were then stained with primary antibodies to LAMP1 for 2 h at 25°C, washed three times in PBS, and incubated with an Alexa Fluor 488-labeled secondary antibody (0.5 µg/mL) for 1 h at 25°C. All cells were then counterstained with Hoechst 33342 (2 µg/mL), rinsed in PBS, mounted in Vectashield (Vector Laboratories Inc., Burlingame, CA), and analyzed by fluorescence microscopy (Nikon Eclipse TE2000S). Western blotting was performed as previously described (84).

## ACKNOWLEDGEMENTS

The authors wish to thank Dr. Dean G. Tang for kindly providing the prostate cancer cell lines, and Drs. Francis Barr, Silvia Corvera, Esteban Dell’Angelica, Bo van Deurs, David Engelberg, Hans-Peter Hauri, Terence Hébert, J. Paul Luzio, Walther Mothes, Suzanne R. Pfeffer, Richard Scheller, Miguel C. Seabra, and Andrew Thorburn for kindly providing expression plasmids. The authors are also grateful to Dr. David Dinsdale (MRC Toxicology Unit, Leicester, UK), Dr. Dwight Romanowicz (ICMB Microscopy & Imaging Facility at UT-Austin), and Pam Whitney (Cell & Tissue Facility Core at MD Anderson Cancer Center) for advice and expertise in TEM sample preparation/analysis and cell sorting. This research was supported by NIH R01 grant R01GM116024 and the University Cancer Foundation *via* the Institutional Research Grant program at the University of Texas MD Anderson Cancer Center (S.B.B.), as well as awards from the Cancer Prevention and Research Institute of Texas (RP160657 and RP180880 to K.N.D.).

## AUTHOR CONTRIBUTIONS

D.J.W., S.V., and S.B.B. conceptualized and designed the experiments. D.J.W., S.V., and D.D. performed the TEM experiments; E.J.C. and K.N.D. performed the *in vitro* PIKfyve kinase assay; M.-D. C. performed endocytosis assays; and D.J.W., S.V., and Z.P. carried out all of the remaining experiments. D.J.W., S.V., S.M., K.N.D., and S.B.B. analyzed and discussed the data; and S.B.B. wrote the manuscript.

## CONFLICT OF INTEREST

The authors declare that they have no conflicts of interest with the contents of this article.

## Notes

### Competing Interest Statement

The authors have declared no competing interest.

