## Supplemental Figure Legends for "Unexpected inhibition of the lipid kinase PIKfyve reveals an epistatic role for p38 MAPKs in endolysosomal fission and volume control"

**fig S1. p38 MAPK inhibitors, but not JNK, ERK, mTOR, or PI3K inhibitors, induce vacuolation.** *(A)* Various transformed and cancer cell lines were treated with DMSO (control), SB203580 (50 μM), or SB202190 (50 μM) for 24 h and assessed for vacuolation by phase contrast microscopy. *(B)* DU145 cells were treated with various kinase inhibitors (50 μM), including SB203580, SP600125, PD98059, rapamycin, and LY294002, and assessed for vacuolation and cell death. *(C)* DU145 cells were treated with SB203580 (50 μM) for 24 h and then washed and replaced with fresh media ± SB203580 for 4-24 h. At each time point following the washout, cells were examined for vacuolation by phase-contrast microscopy (*see* Fig. 1D). *(D,E)* Concentration-dependent inhibition of HSP27 phosphorylation by SB203580 (0 - 100 μM) was determined by western blotting, and individual bands were scanned, quantified with ImageJ software, and plotted as the percent of p-HSP27 inhibited (*see* also Fig. 1E). At each SB203580 concentration, the percentages of vacuolated cells were plotted against the losses in HSP27 phosphorylation and analyzed by linear regression ($R^2$ = 0.82).

**fig S2. SB203580 does not induce the formation of double-bilayered autophagosomes and does not stimulate long-lived protein degradation.** *(A)* HEK-293, HELA, and MEF cell lines were transiently transfected with EGFP-LC3 and treated with SB203580 (50 μM) for 24 h. Some but not all LC3-labeled vacuoles are indicated with arrows (*see* also Fig. 2D). *(B)* ATG5-deficient DU145 cells were treated with SB203580 (50 μM) for 24 h and analyzed by transmission electron microscopy. Representative vacuoles containing partially digested material were magnified (*iv*) 28,000X, (*v*) 36,000X, and (*vi*) 44,000X (bars = 500 and 200 nm). *(C)* DU145, A549, HCT116, and HT-29 cancer cell lines were labeled with [$^{14}$C(U)]-L-Valine

(0.2 μCi/mL media), starved or exposed to SB203580, in the absence or presence of 3-MA, and assayed for LLPD by scintillation counting, as described in the methods. *(D)* Vacuoles induced by SB203580 in autophagy-competent HCT116 cells are enclosed by a single bilayered membrane and contain partially digested material.

**fig S3. SB203580 does not induce vacuolation through dysregulation of early endocytosis.** *(A-C)* DU145 cells were transfected with mCherry-GDI (wild-type or the phosphomimetic S121E mutant); or EGFP-EEA1 (wild-type or the phosphomimetic T1392E mutant). In order to more easily quantitate the number of vacuoles per cell, EEA1-transfected cells were co-transfected with mCherry as it was excluded from vacuoles. Following transfection, all cells were treated with DMSO or SB203580 (50 μM) for 24 h and assessed for vacuolation by fluorescence microscopy (600X). Some but not all vacuoles are indicated with arrows. *(D and E)* DU145 cells were incubated with Dextran-FITC or BSA-FITC for 1 h, in the presence or absence of DMSO (control), SB203580 (100 μM), apilimod (100 nM), or EIPA (100 nM). Cells were then imaged by confocal microscopy and the area of FITC-labeled puncta analyzed using ImageJ software. *(F)* DU145 cells were cotransfected with LAMP1-EGFP and mCherry-Rab9, and then exposed to DMSO (control) or SB203580 (50 μM) for 24 h and examined for colocalization with the vacuoles and/or one another.

**fig S4. SB203580-induced vacuoles label with ER-directed and *cis*-Golgi markers, but not *medial* or *trans*-Golgi markers.** *(A,B)* DU145 cells were transiently transfected with ER-targeted EGFP-Cyt. b5, ERGIC-EGFP, *cis*-Golgi markers GRASP65-EGFP or EGFP-GOS28, *medial*-Golgi marker GRASP55-EGFP, or *trans*-Golgi marker TGN38-EYFP. Cells were then treated with DMSO (control) or SB203580 (50 μM) for 24 h and examined by fluorescence microscopy. Some but not all vacuoles are indicated with arrows. TGN38-EYFP labeled smaller vesicles in some SB203580-treated cells but was not observed on the larger vacuoles. *(C)* DU145 cells, cotransfected with the *cis*-Golgi marker GRASP65-EGFP and lysosome marker mCherry-CD63, were exposed to DMSO (control) or SB203580 (50 μM) for 24 h and examined by fluorescence microscopy.

**fig S5. SB203580-induced vacuolation is reversible and results from an osmotic imbalance.** *(A)* Representative images for Fig. 5E. *(B)* DU145 cells were treated with SB203580 (50 μM) for 24 h and then washed and replaced with fresh media ± SB203580 (50 μM) for 4-24 h. At each time point following the washout, cells were stained with LysoTracker$^{TM}$ Green and analyzed by flow cytometry for acidification. *(C)* Following a 3 h washout ± SB203580 (50 μM), changes in vacuolation were examined by immunofluorescence microscopy using an anti-LAMP1 antibody (panels *i,ii*: 600X) and by transmission electron microscopy (panel *iii*: 4,400X, bar = 2 μm; panel *iv*: 28,000X, bar = 500 nm). Following a 1 h washout of SB203580, the large cytoplasmic vacuoles began to collapse, flatten out, and undergo fission (panel *v*: 5,600X, bar = 2 μm; panel *vi*: 14,000X, bar = 500 nm). *(D)* DU145 cells were treated with SB203580 (50 μM) for 24 h, in the presence or absence of sorbitol (1 mM), and evaluated for vacuolation by phase-contrast microscopy. *(E)* DU145 cells were exposed to SB203580 (50 μM) for 24 h, followed by the addition of DMSO or Bafilomycin A1 (125 nM) for an additional 24 h. Vacuolation was then evaluated by phase-contrast microscopy and quantified using ImageJ software. *(F)* Cartoon illustrating various ion channels that maintain normal osmotic balance in LELs. *(G-J)* DU145 cells were treated with SB203580 (50 μM) for 24 h, in the presence or absence of a Na+-H+

exchanger inhibitor (Zonaporide, 25-500 μM), a CLC chloride channel inhibitor (4,4′-Diisothiocyanatostilbene-2,2′-disulfonic acid, *i.e.* DIDS, 1-4 mM), a CFTR chloride channel inhibitor (CFTR inhibitor-172, 5-75 μM), or an NCX $Na^+/Ca^{2+}$channel inhibitor (SN-6, 10-200 μM) The cells were then evaluated for vacuole formation by flow cytometry (LysoTracker™ Green) or phase-contrast microscopy.

**fig S6. SB203580 and SB202190 inhibit the production of PI(3,5)P2 and induce vacuolation similar to established PIKfyve inhibitors.** *(A)* HCT116 cells were treated with YM201636 (1 μM) for 24 h and analyzed by transmission electron microscopy. *(B)* DU145 cells were treated with SB202190 (50 μM), SB203580 (50 μM), YM201636 (1 μM), or apilimod (50 nM) for 24 h and evaluated for PI(3,5)P2 levels by immunofluorescence microscopy. *(C)* DU145 cells were treated with SB203580 (50 μM), SB202190 (50 μM), BIRB-796 (50 μM), YM201636 (1 μM), or apilimod (50 nM) for 24 h and then immunoblotted for HSP27, p-HSP27, and β-tubulin.

**fig S7. The p38 MAPK inhibitor BIRB-796 enhances vacuolation induced by PIKfyve inhibitors.** *(A-C)* Wild-type DU145 cells were treated with YM201636 (500 nM) or apilimod (20 nM), in the presence or absence of BIRB-796 (50 μM) for 24 h and evaluated for vacuolation by phase-contrast microscopy and ImageJ analysis software. *(D)* Representative images for washout experiments; see legend for Fig. 7H for further details.
