## Supplementary figures and images for "Unexpected inhibition of the lipid kinase PIKfyve reveals an epistatic role for p38 MAPKs in endolysosomal fission and volume control"

### Supplemental Figures 1-7

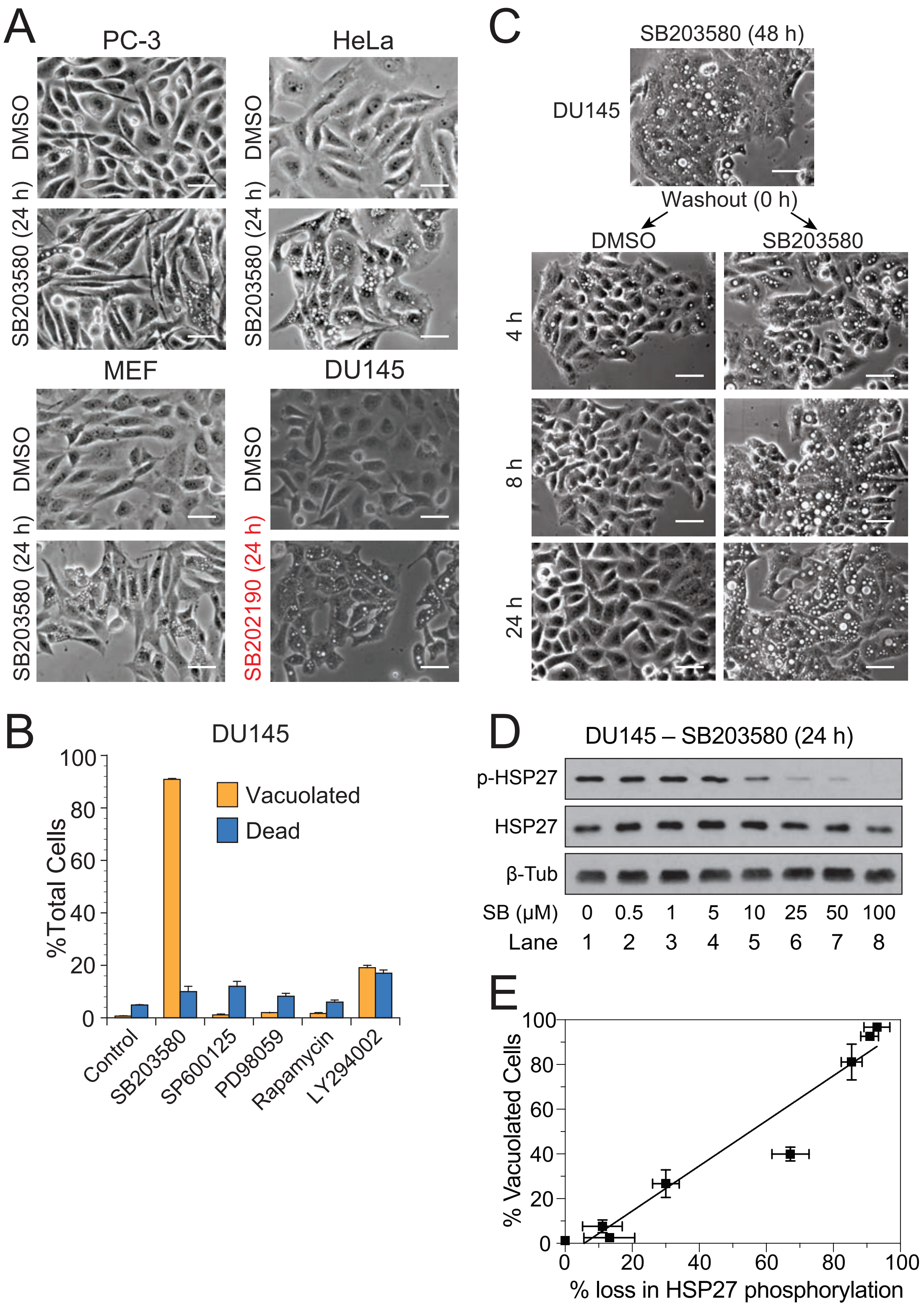

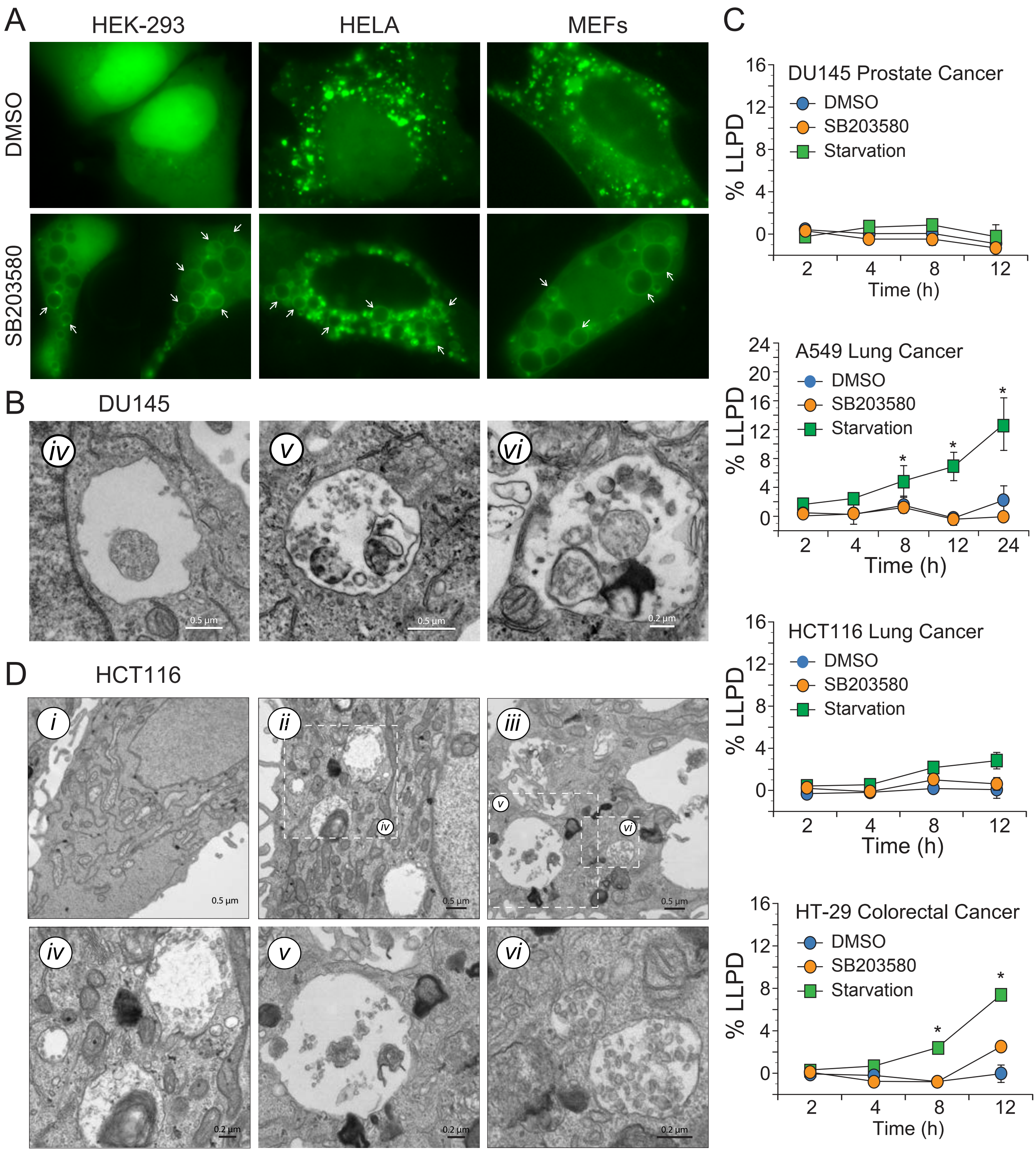

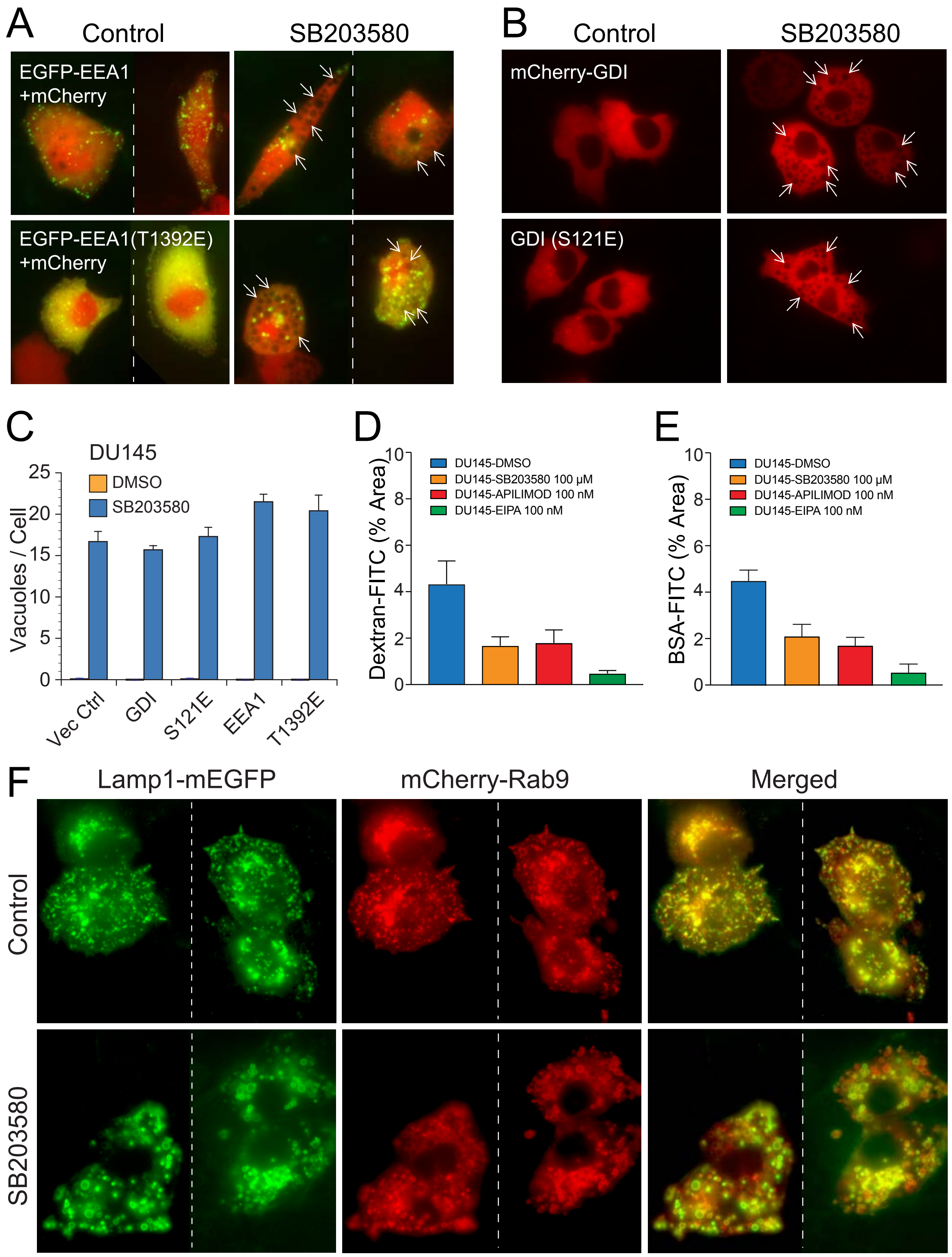

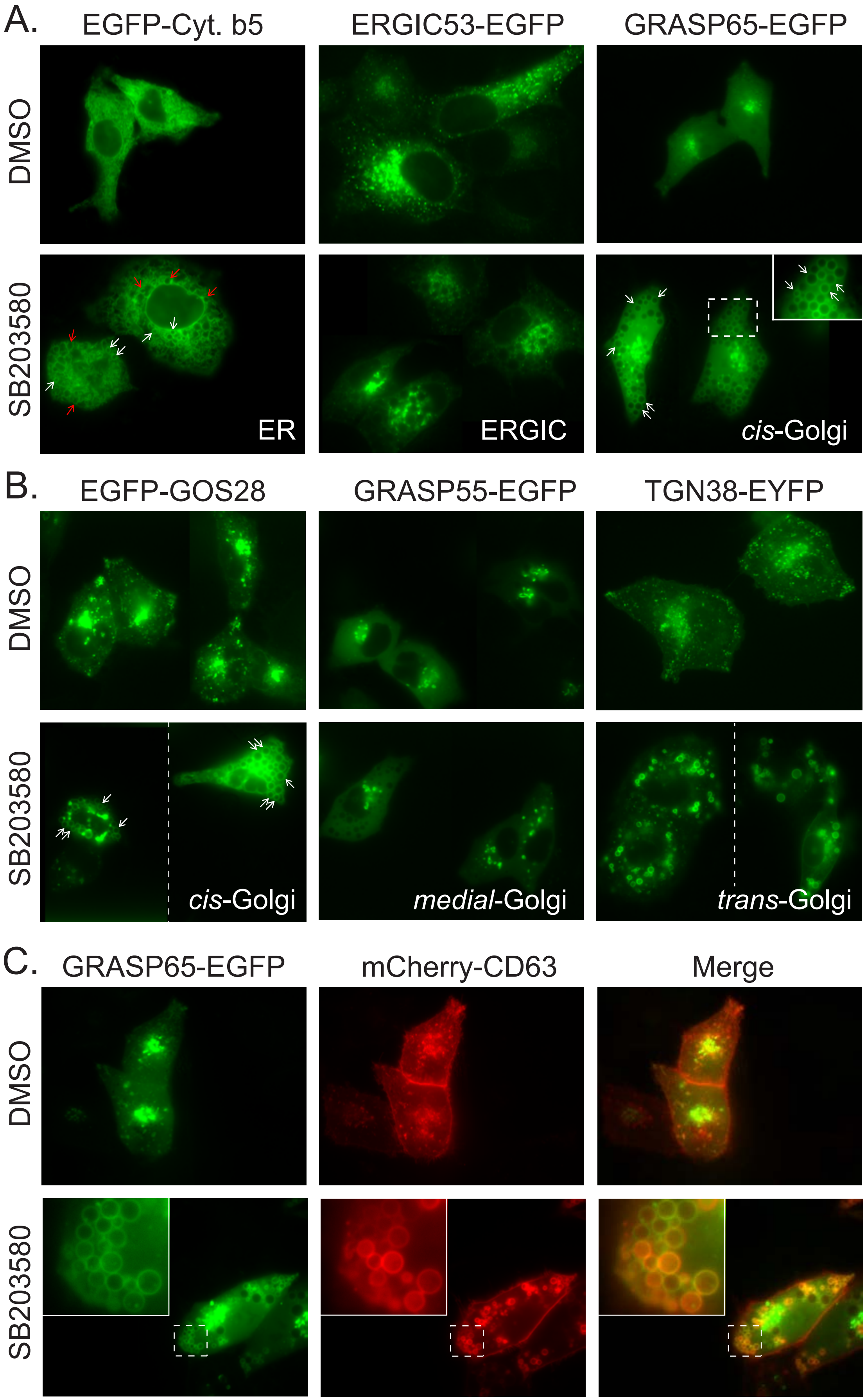

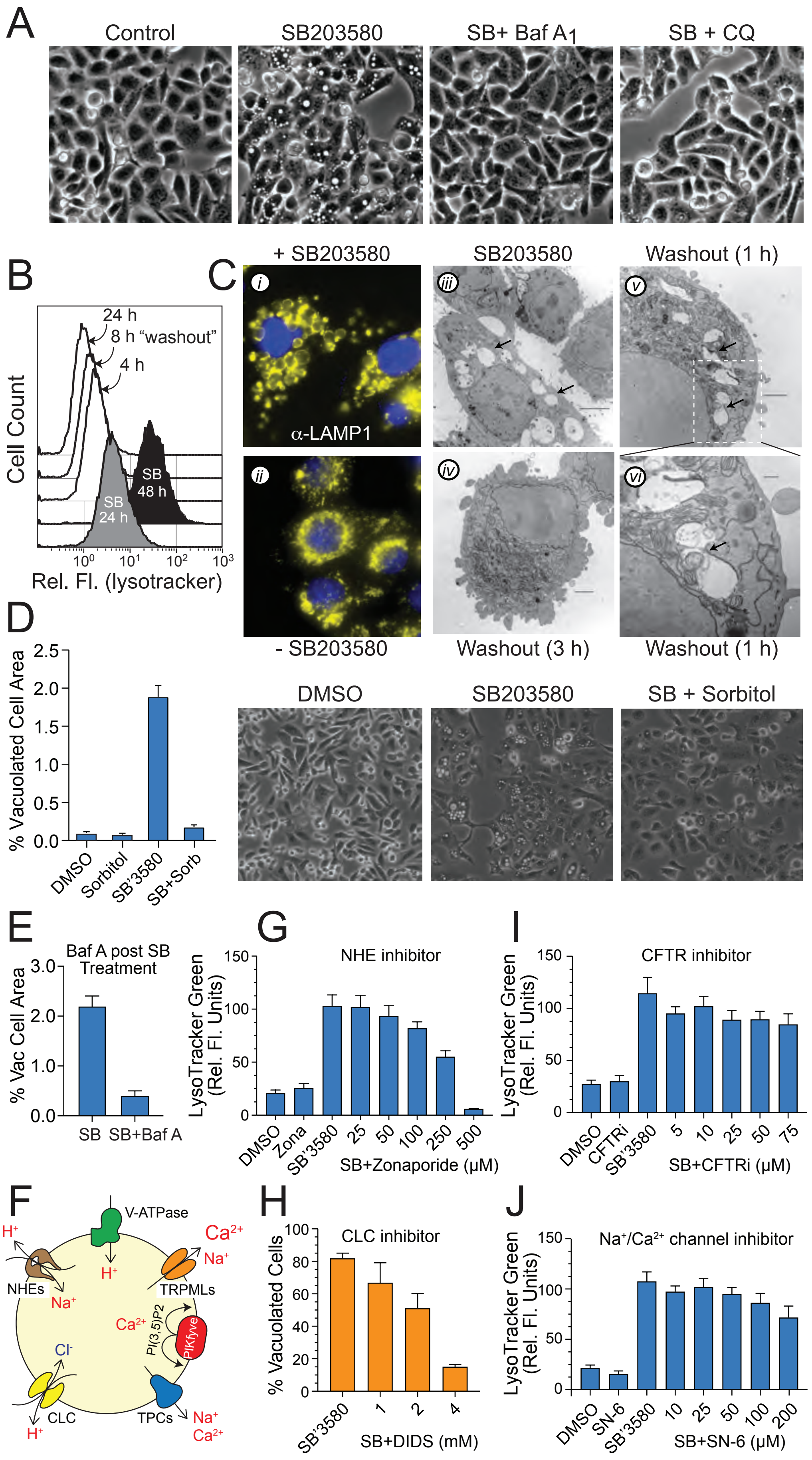

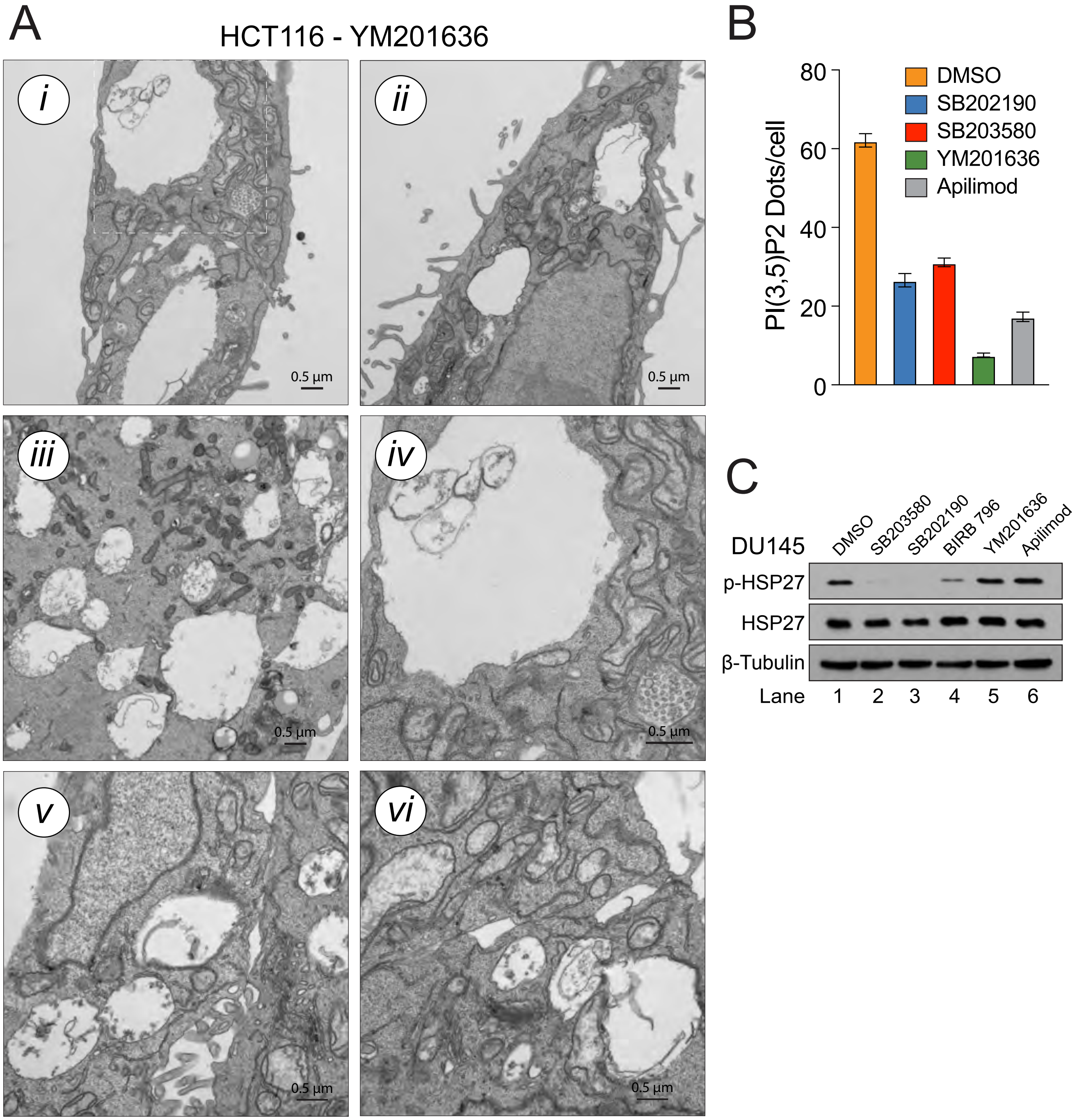

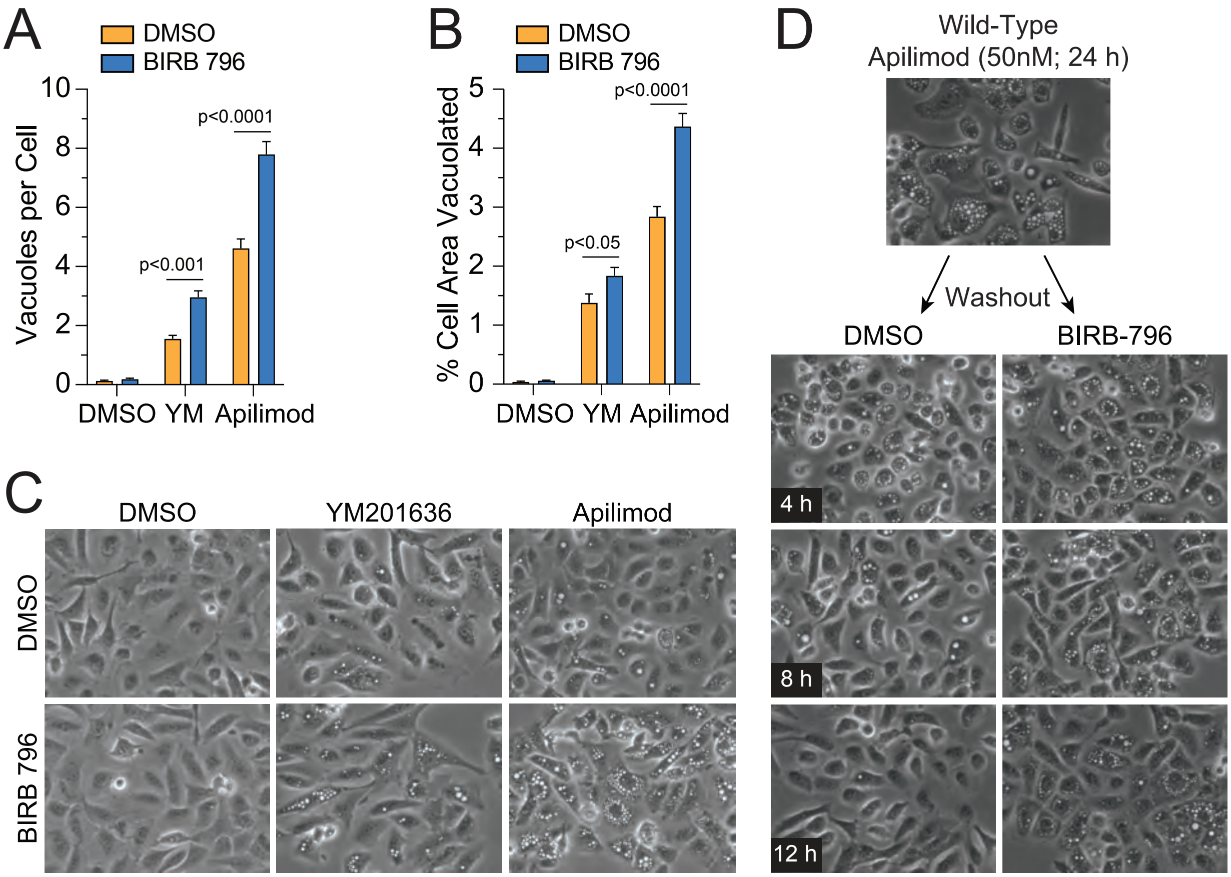
